# Inference of drug off-target effects on cellular signaling using Interactome-Based Deep Learning

**DOI:** 10.1101/2023.10.08.561429

**Authors:** Nikolaos Meimetis, Douglas A. Lauffenburger, Avlant Nilsson

## Abstract

Many diseases emerge from dysregulated cellular signaling, and drugs are often designed to target specific nodes in cellular networks e.g. signaling proteins, or transcription factors. However, off-target effects are common and may ultimately result in failed clinical trials. Computational modeling of the cell’s transcriptional response to drugs could improve our understanding of their mechanisms of action. Here we develop such an approach based on ensembles of artificial neural networks, that simultaneously infer drug-target interactions and their downstream effects on intracellular signaling. Applied to gene expression data from different cell lines, it outperforms basic machine learning approaches in predicting transcription factors’ activity, while recovering most known drug-target interactions and inferring many new, which we validate in an independent dataset. As a case study, we explore the inferred interactions of the drug Lestaurtinib and its effects on downstream signaling. Beyond its intended target FLT3 the model predicts an inhibition of CDK2 that enhances downregulation of the cell cycle-critical transcription factor FOXM1, corroborating literature findings. Our approach can therefore enhance our understanding of drug signaling for therapeutic design.

## Introduction

In many diseases, such as cancer, alterations in gene expression or protein function lead to dysregulated intracellular signaling, with pathological effects^1–4^. This may be counteracted by perturbing cellular signaling using drugs, in particular small molecules have been used for decades to revert cells to a healthy state or kill cancerous cells^1^, e.g. inhibition of Ras-mediated signaling in anticancer therapy^5^. This approach aims to affect signaling through specific drug-target interactions, but the drugs do not necessarily function through their proposed mechanism of action (MoA)^6^ and off-target effects are common^7^. Understanding the contributions of on- and off-target effects of drugs is important for the development of safe therapeutics and their success in the clinic.

Systems pharmacology approaches have been developed to decipher the MoA of drugs. Several of these utilize data from chemical perturbation experiments^8–10^. For example, these approaches may utilize the transcriptomic profiles of perturbed cells to identify key genes associated with specific therapeutic or adverse effects^11^ or elucidate their signaling mechanism based on their gene expression profile and large datasets of known drug-target interactions^12,13^. With the advent of Machine Learning (ML) and large-scale high throughput screening (HTS) datasets, such as the L1000 dataset^14^, consisting of thousands of drug perturbations tested on cancer cell lines, these approaches have become more efficient, e.g. leading to the identification of novel potential therapeutic targets^15^ and to direct characterization of the transcriptomic profile of perturbations^16^. However, these approaches do not explicitly model the signal propagation that underlies these effects and their predictions can therefore not be directly interpreted in terms of molecular mechanisms.

Signaling networks provide a scaffold to comprehensively describe a drug’s MoA. Molecular networks have been used to agglomerate signature MoA predictions^17^, as the basis for large-scale computer models to facilitate genome-scale simulations of perturbations^18,19^. This has become feasible due to the extensive characterization of the intracellular signaling network^20,21^ and improvements in parameter fitting methods. For example, in early work, Saez-Rodriguez and Alexopoulos et al.^22^, used Boolean modeling on a small small-scale signaling network to predict inflammatory signaling in HEPG2 cell lines while inferring interactions that were missing from the initial network. In more recent work, Fröhlich et al.^23^ developed a large-scale mechanistic model using ordinary differential equations (ODEs) to predict the response to drug perturbations in 120 different cell lines. Alongside the signal network, this model relied on a sparse network of drug-signaling protein interactions that was manually curated from the literature. However, despite major advances, the parameter fitting of ODE-based models could require problematically long computational times when applied to genome-scale networks.

Artificial neural networks (ANNs) allow for rapid parametrization of large-scale models. While the interpretability of such models could be a concern, this can be circumvented by constraining them to only allow mechanistically plausible predictions. For example by only allowing connections in the ANN that correspond to interactions of the intracellular signaling network, which has been used to predict receptor stimulation from gene expression data^24^. We recently expanded on this concept and developed a framework to simulate signal propagation using a recurrent neural network (RNN), thereby allowing feedback loops to be incorporated in the formalism^25^. Using this framework, termed, Large-scale knowledge EMBedded Artificial Signaling network (LEMBAS), we successfully trained a model to predict the transcription factor activity of macrophages in response to different ligand stimulations. For this, we took advantage of both a prior knowledge network of signal transduction, and a transcriptional regulatory network^26^. The latter was used to infer transcription factor activity from gene expression using the VIPER algorithm^27^, which tests for regulon enrichment on gene expression signatures. We also adapted the LEMBAS framework to replicate the prediction of drug responses from the Fröhlich study with indistinguishable accuracy and much faster parametrization time. However, both of these approaches depend on prior knowledge of drug-target interactions and were not designed to infer new drug targets. Because it is improbable that all drug-target interactions have already been discovered, in particular for newly developed drugs, inference of new interactions could be of importance to completely explain the effects of drugs. Many different ML approaches have been developed to infer new potential interactions using bioactivity data, dose responses, and large databases of prior knowledge containing known drug-target interactions^15,28–30^. However, current ML approaches focus on inferring single drug-target interactions or binding affinities, either based on chemical structures^31,32^ or gene expression profiles^15^, without fully utilizing the signaling network. They thus lack direct interpretability and the ability to comprehensively describe the signaling cascades arising from off-target MoA.

Here we have developed an approach to predict network-wide signaling responses to drugs that considers both on- and off-target effects. We expand the ANN-based signaling framework^25^ to combine a prior knowledge network of signaling^20^, a network of known drug-target interactions, and the drugs’ chemical structure similarity with other drugs, to simultaneously infer drug-target interactions and simulate the regulatory effect of known and inferred interactions in drug perturbation experiments. We use publicly available data on the transcriptomic response to drug perturbations, that we process further to infer transcription factor activities. We use the data to train cell-line-specific signaling models that we use to identify potential off-target effects of drugs alongside MoAs that can explain them. We validate the inferred interactions using an independent dataset and explore some of the predicted MoAs using in silico simulations and public gene knock-out data.

## Results

### A model for predicting network-wide signaling of drugs via modeling of on- and off-target effects

We developed an approach (denoted as DT-LEMBAS) for predicting the regulatory effect of drug perturbations, while simultaneously inferring unknown drug-target interactions (Figure 1A). The model consists of two interconnected sub-modules: the first module takes drugs’ concentration as input and gives their signaling effect on drug targets as output, while the second module is LEMBAS, a published model of intracellular signaling that takes a drug’s signaling effects as input and returns the transcription factor (TF) activity as output^25^. LEMBAS is a recurrent ANN model of intracellular signaling, where the connections are based on prior knowledge of the intracellular signaling network, thereby constraining the model to mechanistically plausible predictions.

**Figure 1:**
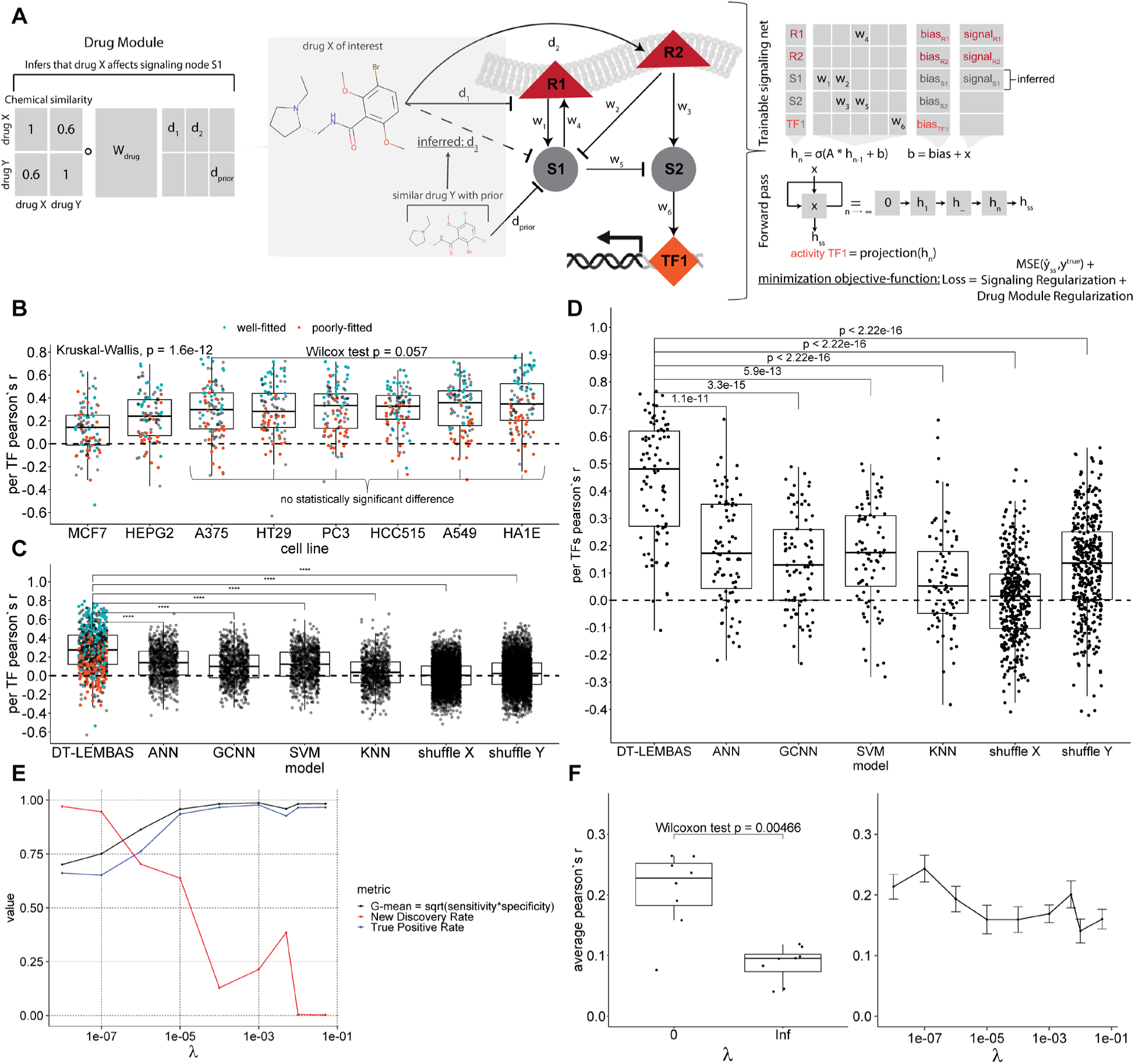
Model architecture and basic performance metrics. **A)** DT-LEMBAS’ model architecture, consisting of two interconnected sub-modules: 1) a drug module that calculates drug signaling via drug-target interaction inference, based on known drug-target interactions and chemical similarity with other drugs and 2) a LEMBAS-based recurrent ANN, modeling intracellular signaling. **B)** Performance of our approach in different cell lines. **C)** Performance comparison in validation sets with standard machine learning approaches, for every TF in every cell line, using Pearson’s r between predicted and actual TF activity. The model is also compared to two randomized models, derived by shuffling the inputs (X) and outputs (Y) on training. **D)** Performance comparison in validation sets, by looking only at the TFs well-fitted (here top 10%) during training. **E)** The optimal geometric mean of sensitivity and specificity for inferring drug-target interactions, and the NDR and TPR for the same gradient cut-off, at different levels of regularization **F)** Average performance across all TFs in every cell line for different levels of regularization. A two-sided unpaired Wilcoxon test was used to compare “*infinite”* and “*0”* regularizations.

In the case of drug perturbations, such as treatment with small molecules, the prior knowledge of the drugs’ molecular interactions may be incomplete, thus creating the need to infer potential off-target signals. To achieve this we utilize both known drug-target interaction information, taken from the Broad’s Institute Repurposing Hub^33^, and pre-calculated chemical similarity between drugs, using their ECFP4 molecular fingerprints^34^. We encode the drug-target interactions as a trainable spare weight matrix and the chemical similarity as a drug-drug similarity matrix, forming a pre-defined drug/target space (Figure 1A). The concentration of a drug of interest is taken as input and, based on the similarity with other drugs and its known targets, the module is allowed to infer potential drug-target interactions that are propagated as input signal to the LEMBAS module. We train the combined model to fit TF activity data while minimizing a few regularization terms, aimed at controlling the number of new inferred drug-target interactions from the drug module, alongside regularizations and other priors previously developed for LEMBAS^25^ (see methods).

### Performance in predicting activities of individual transcription factors

To train our model, and evaluate its performance in predicting activities of individual transcription factors, we used gene expression data from the L1000 dataset^14^. As the purpose of this study is to examine the short-term signaling of drugs, to avoid self-regulatory effects we excluded long-period experiments and used only perturbations where cell lines were treated with a drug for less than 12 hours.

As ANNs are known to overfit the data, before performing any subsequent downstream analysis of the predicted MoA of drug perturbations it is useful to determine which TFs the model is able to predict correctly. Similarly, it is necessary to ensure that the predictions of the drug module generalize sufficiently well for the inferred drug-target interactions to be trusted. Cross-validation is a common validation strategy, where some of the data is withheld from the training set, however, in this dataset, drugs only appear once per cell line, meaning that there would be no training data available for the drug using this approach. An additional challenge for ML methods using chemical representations is misleading high performance due to memorization of similar structures^35^.

To circumvent these issues, we devised a validation strategy (Supplementary Figure 1) that makes use of data from different cell lines to construct the drug module, while applying a cross-validation schema to the signaling module. Specifically, we select cell lines with data from at least 200 drugs in common, resulting in 9 cell lines. We then use the cell line with the most samples available (VCAP) to train a model. For the remaining 8 cell lines we keep the drug module unchanged and train only the signaling part of the model using 80% of the drugs available while validating using the remaining 20%, i.e. fivefold cross-validation. We confirm that validation drugs are highly dissimilar in their chemical structure from training drugs, thereby avoiding information leakage from similar drugs (Supplementary Figure 3). We find that the model predicts the activities of many TFs with high accuracy, and a similar behavior is observed across all cell lines (Figure 1B). Notably, there is a big variation in the performance between individual TFs, and since we are aiming to utilize some of them for downstream analysis it would be useful to find a principled way to identify high-performing TFs. We hypothesize that some TFs will be poorly fitted by the model during training due to various reasons, e.g. because their data may be noisy, that they may not be contributing to the transcriptomic profile of the cell, or that their activity cannot be explained by the prior knowledge signaling network. Indeed, TFs that are well-fitted during training (rank≤25% in training based on Pearson correlation) are overrepresented among the high-performing TFs in validation, while most TFs that are poorly fitted (rank≥75% in training) also perform poorly in validation (Figure 1C, Supplementary Figure 6). Similar results can be observed if the drug module is initially trained in other cell lines such as A375 and A549 (Supplementary Figure 5).

To determine if these validation results are in line with what could be expected given the data, we benchmark them against four basic machine learning techniques. Specifically, we compare the models’ ability to predict TF activities with: 1) an ensemble of 50 simple feed-forward artificial neural networks (ANN) that take as input the ECFP4 molecular fingerprints of drugs 2) an ensemble of 50 Graph Convolutional Neural Networks (GCNN)^36^ representing drugs’ chemical structures as graphs, 3) an ensemble of 50 Support Vector Machines (SVMs), and 4) an ensemble of 50 KNN models. Additionally, as two types of null models, we train two models where we 1) shuffle the input matrix of drug concentrations (X) during the training of the drug module, thereby generating a randomized drug module, or 2) we shuffle the outputs (Y) during re-training of the signaling part for each cell line, generating a randomly weighted signaling network. Our model outperforms these approaches, as well as the null models, based on a non-parametric two-sample Wilcoxon test (Figure 1C). The validation performance for the top 10% fitted TFs during training is generally high (Figure 1D), achieving an average Pearson correlation of ∼0.5 (Supplementary Figure 2B), with the performance of some TFs higher than ∼0.8 and p-values ≤ 10^−6^ (see Supplementary Figure 4 for the adjusted p-values for all of the correlations). This suggests that we can rely on the predictions for some of the TFs in our subsequent analysis.

### Constraining the number of inferred interactions via weight regularization

We make use of the assumption that drugs will not interact with most targets to make more specific predictions. To control the number of inferred interactions, we utilized an L2-based regularization scheme for the weights of the drug module such that infinite regularization constrains the module to only make use of known drug-target interactions and zero regularization allows every possible interaction without penalty (see methods). Since we cannot know in advance which targets a drug does not affect (true negatives) we instead aim to find a good trade-off between sensitivity and specificity in inferring interactions, as well as prediction performance. We utilized an integrated gradient score approach^37^ (see methods) to quantify the confidence in a drug affecting a target node in the signaling network and we inferred interactions by identifying a cut-off for the absolute value of that score (see methods). With increasing regularization, the trade-off between sensitivity and specificity saturates (Figure 1E) when inspecting their geometric mean (optimal G-mean) at the cut-off that maximizes it. To quantify the amount of interactions at different regularization levels we define a metric, New Discovery Rate (NDR), as the number of new interactions inferred divided by the number of total interactions inferred by the model. We find that for increasing regularization levels, this metric decreases and slowly goes to zero, as intended (Figure 1E). Meanwhile, for increasing regularization levels the True Positive Rate (TPR) increases and saturates, indicating that with increasing regularization the model depends more on prior knowledge, and as intended it does not exclude a lot of prior knowledge interactions to reduce the total inferred interactions (Figure 1E). Similarly, for every regularization level at different gradient score thresholds the G-mean and TPR increase and start saturating after λ= 1E-04 while the NDR decreases until it becomes almost zero (Supplementary Figure 7A). This result appears to be robust to using a different error-based method to infer interactions (see methods, and Supplementary Figure 7B). Finally, for the average performance of individual models (not the ensemble) trained using different regularization levels, we observe that zero regularization outperforms infinite regularization (Figure 1F). This indicates that the addition of inferred interactions contributes to the model’s predictive power. However, there is not a clear trend for intermediate levels of regularization, nevertheless, it seems that for the regularization level λ= 5E-03, the performance is slightly higher than its neighboring levels (Figure 1F). This could perhaps be due to the locally higher NDR at that regularization level (Figure 1E and Supplementary Figures 7). Because of this, alongside the higher performance, we selected this regularization level for the models trained in this study (including the models in Figure 1C).

### Inferring drug-target interactions with integrated gradient scores

The model is constructed to allow inference interactions that are not part of the prior knowledge to better explain transcriptional data. To extract which drug-target interactions have been inferred, we use integrated gradients to assign an importance score to each interaction (see methods). In the case of a linear drug module, used in this study, the score is proportional to the module’s weights (Supplementary Figure 8). A negative score corresponds to a potential inhibition of a target node from a drug of interest, while a positive score corresponds to activation.

To identify a cut-off level for the score we investigate how the model’s performance decreases as more interactions are removed (see methods). Briefly, for each drug, we successively remove more interactions, based on the absolute value of the score, and determine how this affects the error of the model in predicting the activity of all TFs (Figure 2A). As can be expected, removing interactions with low scores does not affect the error of the model, while at some critical level, the error sharply increases and plateaus. This means that the model’s low-scoring interactions are not needed to explain the TF activity, while for high-scoring interactions, the error of the trained model increases dramatically. For each drug in each trained model, we define the cut-off at a 25% percentage increase in error. This approach was chosen because of its high NDR (Supplementary Figure 7) and because it allows us to infer many new interactions which at the same time are necessary for the model to correctly predict the TF activity. Subsequently, we utilize the ensemble of models to also score the confidence in inferring an interaction by using the frequency of appearance in multiple models.

**Figure 2:**
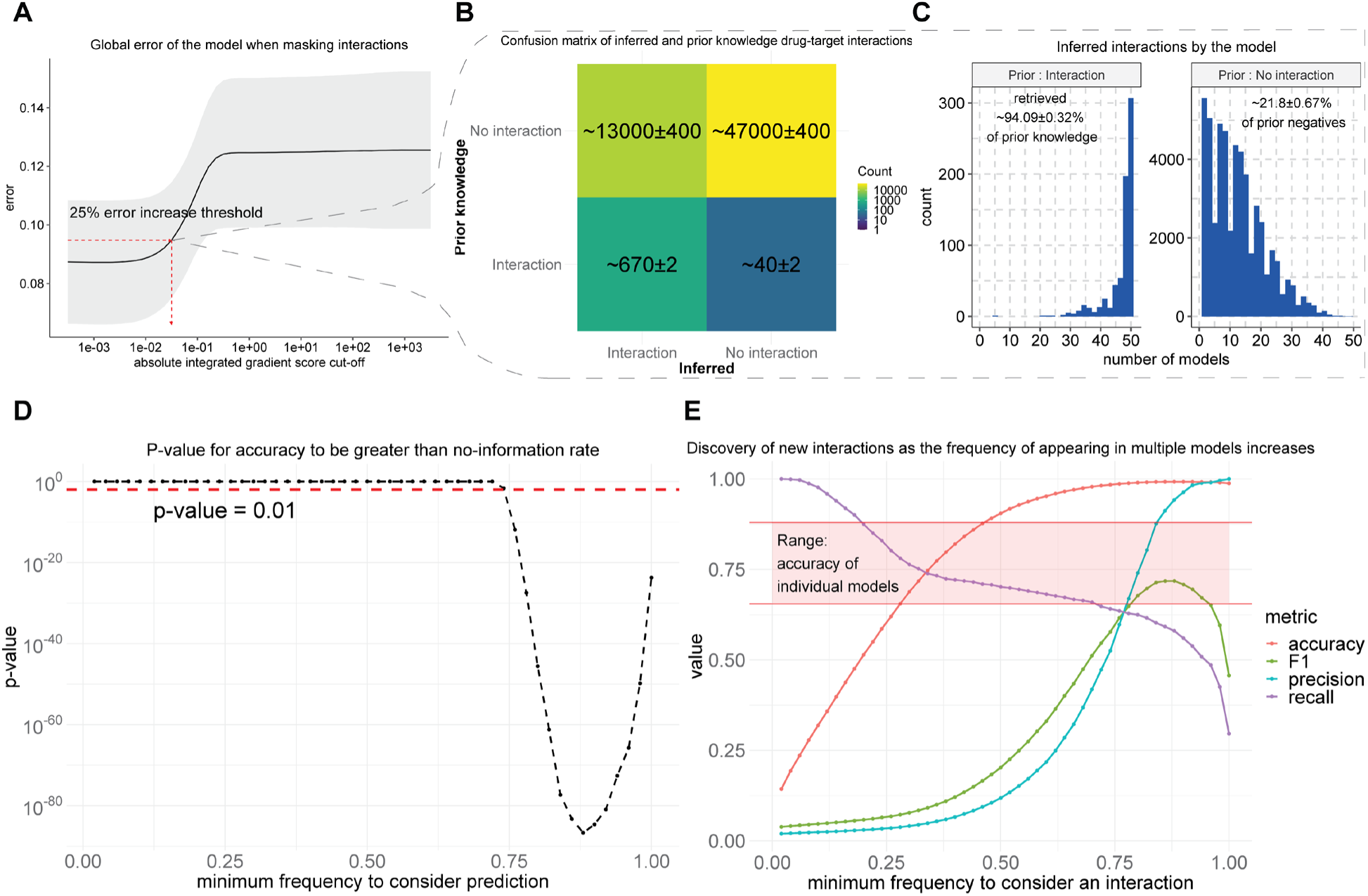
Inferring drug-target interactions in the A375 cell line from the drug module. **A)** Error of the model as more important drug-target interactions, according to their integrated gradient score, are removed. **B)** Average confusion matrix from 50 trained models for the inferred drug-target interactions. **C)** Percentage of prior knowledge drug-target interactions and previously unknown interactions retrieved, and their corresponding frequency of appearance in multiple models. **D)** Classification performance of our approach by considering as ground truth the interactions contained both in the Broad’s Institute Repurposing Hub^33^ and in DrugBank^13^. Performance is calculated for an increasing frequency of an interaction appearing in multiple models. **E)** P-values from comparing accuracy with the accuracy obtained by assigning everything to the predominant class (No Information Rate: NIR), for multiple frequency scores, in this imbalanced dataset where most drugs do not interact with most targets.

The first step to evaluating the validity of this approach to infer drug-target interactions is whether it can retrieve most of the prior knowledge interactions (on-target effects), as these are expected to be able to explain at a large level the observed transcriptional profile. Indeed, when training a model for the A375 cell line, we can retrieve most of the interactions in the prior knowledge used in training the model while also inferring approximately 13000 more interactions (Figure 2B-2C), which can potentially be undiscovered direct drug-target interactions, indirect effects or false interactions (false positives). It seems that prior knowledge interactions are inferred by most of the models in the ensembles, while undiscovered interactions appear mostly with low frequency, with some of them appearing in many models (Figure 2C). We observe similar results when training models and inferring drug-target interactions using the A549 and VCAP cell lines (Supplementary Figure 9). Based on this, we hypothesized that it could be possible to predict if an inferred interaction is a true direct interaction, based on the number of times it was inferred by different models.

### Evaluating the inference of direct interactions by an ensemble of models

We make use of an independent drug-target interaction database to evaluate the predictive power of the model. While we know the existing true drug-target interactions (true positives), the true negatives are unknown, and it is not clear to which extent predicted interactions can be trusted. To partially overcome this limitation, we make use of a more comprehensive database, DrugBank^13^, for the drugs present in our trained framework, to establish a set of true interactions that were not present in the prior knowledge used to construct the model. We then attempt to predict these interactions depending on how frequently they are inferred in our models. From a frequency of 0.75 (interaction inferred for 37 out of 50 models), there is a statistically significant difference between the model’s accuracy in predicting drug-target interactions and the null accuracy obtained by assigning everything to the predominant class (No Information Rate: NIR) (Figure 2D). In addition to accuracy, we also consider the following evaluation metrics: precision, recall, and the F1 score, which is the trade-off of precision and recall for imbalanced data (Figure 2E).

We find that including interactions that appear in any model is much too lenient, resulting in poor precision (1.97%) and accuracy (14.35%), and the predictions of individual models do not perform better than chance with an accuracy of ∼75%. However, for increasingly frequent interactions both precision and accuracy increase markedly, with perfect precision (100%) for the most frequent predictions (Figure 2E). At high inference frequencies recall decreases drastically, meaning that the inference threshold may be too strict, The F1 score (which is a trade-off between precision and recall) generally increases until it reaches a maximum of 71.84 % for an interaction appearing with a frequency score of 0.88 (44 out of 50 models). This is the same number of models which corresponds to the higher accuracy (99.22%) and the lowest p-value signifying statistical significance in the difference between accuracy (99.22%) and NIR. We observe similar results for models trained on A549 and VCAP cell lines (Supplementary Figure 10). This evaluation showed a higher accuracy than we could expect from naively guessing that an interaction does not exist, which supports the hypothesis that interactions appearing in multiple models are more likely to correspond to direct interactions which enables the potential for inferring novel drug-target interactions. We provide all inferred drug-target interactions alongside frequency scores in Supplementary Data File 1, e.g. Dacinostat, a known histone deacetylase inhibitor, is found in both A375, A549 and VCAP cell lines, by more than 40 models, to interact with KDR which is a type III receptor tyrosine kinase.

We tested if the inferred off-target effects could help explain the lethality of drugs. Off-target effects are primarily thought to cause side effects, but instead, they may contribute to the drug’s efficacy in some cases^8,38^. We tested if the inferred targets could help predict the lethality of drugs tested on different the 9 cell lines in our study that were also present in the NCI60 drug screen^39^. Inspired by Vijay & Gujral who developed an ANN model to predict changes in cell migration of cancer cells using drugs’ target profile^40^, we trained random forest (RF) models to predict lethality using the drug targets and cell line identity as input and keeping data from experiments on the A549 cell line as an independent test set. However, the performance of an RF model trained on the prior knowledge data and the one trained on inferred interactions (with a 0.88 frequency score) performed equally well (Supplementary Figure 13). Both RF models achieve a correlation of 0.94 in the test set (p-values 1.5×10^-6^ and 2×10^-6^ respectively). It seems for the data used here the on-target effects can already fully explain the lethality observed, and that the inferred interactions do not contribute further to the performance. This performance is also dependent on the number of models used to infer interactions (Supplementary Figure 12).

### Identification of transcription factors regulated by off-target effects

After establishing frequency thresholds for trusting predicted drug-target interactions, we make use of the signaling module to investigate their predicted mechanism of action (MoA), in terms of inducing TF activity. We first identify whether the model predicts that there are marked off-target effects in response to a perturbation, by removing all of the input signal outputted by the drug module except the signal corresponding to the known targets and use it as input to predict the induced TF activities (Figure 3A). We consider the difference between the models’ original predicted TF activities and the ones where off-targets are masked out (*ΔTF*) as a proxy for the magnitude of the off-target effects on specific TFs. Samples where a TF has been activated (here considering activity≥0.75) or inhibited (activity≤0.25), and with a high off-target effect are of interest for investigation (Figure 3B). When this contributes to the observed direction of TF regulation, it may be considered a perturbation with off-target effects. We further restrict our analysis to TFs whose activity is predicted well by the model (of the A375 cell line), by making sure that the average of the performance in validation and training of the model for that TF is higher than 0.6. An example of this is the case of the drug Lestaurtinib, where the model predicts an inhibitory off-target effect on FOXM1 (Figure 3B). FOXM1 is a TF critically associated with the cell cycle, considered a master regulator overexpressed in most human cancers^41^. For this reason, we select FOXM1 and Lestaurtinib for further analysis, but more drugs that have an off-target effect on some TFs, in A375, A549 and VCAP cell lines, are provided in Supplementary Data File 2, together with their activity, off-target and performance score.

**Figure 3:**
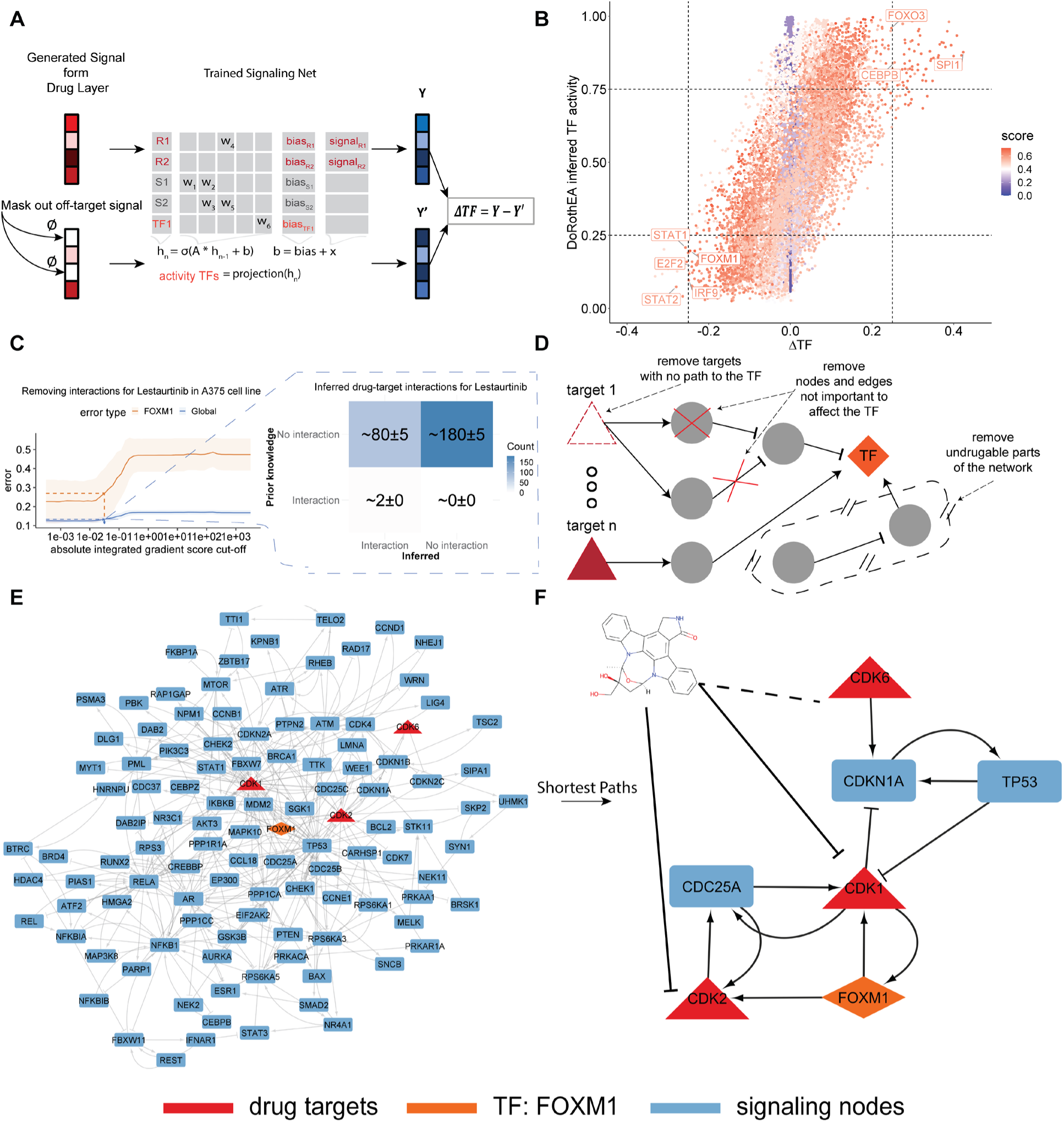
Process for interpreting off-target effects, with a case study for Lestaurtinib’s effects on FOXM1. **A)** The difference between the model’s predicted TF activity and the activity if the off-target signal is removed is used as a measure for the off-target effects on specific TFs. **B)** The activity of the TFs compared with the predicted off-target effects alongside a confidence score from the average performance in training and validation. **C)** Inferring the drug-target interactions using multiple models and the global error approach previously discussed. Here the example of Lestaurtinib is shown. **D)** The whole signaling network is trimmed by removing unimportant edges and nodes to control the TF of interest, stopping the process when there is no path from the inferred targets to the TF of interest. This process is repeated for every trained model and only frequently appearing edges and nodes are kept. **E)** The trimmed ensemble network explaining the off-target effect that leads Lestaurtinib to inhibit more FOXM1, via previously unknown drug-target interactions. **F)** A version of the trimmed network that only considers the simplest paths that connect every target to FOXM1. Lestaurtinib, according to the model inhibits CDK1 and/or CDK2 kinases, while it interacts with an uncertain sign with CDK6.

### A subnetwork explaining off-target effects of Lestaurtinib

We infer all off-target interactions for the drug Lestaurtinib. As previously described (Figure 2A), we infer drug-target interactions by progressively masking potential interactions based on their integrated gradient score and calculating the error of the models for predicting the activity of TFs, until a sharp increase in the error appears. We repeat this process just for Lestaurtinib inspecting the effects on each TF independently. Strikingly the error of FOXM1 follows a trend similar to the average error across all TFs (Figure 3C). Using this approach, we identify a cut-off for the gradient score and infer on average 82 potential interactions out of the 259 available in our training space per model (Figure 3C). Two targets of Lestaurtinib exist in the prior knowledge used to train the model: NTRK1 and FLT3, and both of them are retrieved for all of the models (50 of 50). Additionally, while not in the prior knowledge used for training, DrugBank has another two targets for Lestaurtinib in our target space: EGFR and ADRB1, these are inferred in 16 and 27 of the 50 models respectively. This means that the model retrieves the prior knowledge most of the time, and additionally infers many other interactions not in the training set that act to explain off-target effects.

We extract a sub-network explaining the MoA effects of Lestaurtinib. After inferring new targets and identifying a TF with prominent off-target effects, we use the model to construct a smaller signaling network explaining the mechanism of action for the off-target effects. For this we remove nodes and edges in the trained signaling network models that are not important for regulating the activity of the TF (Figure 3D). This is based on an importance score (see methods) where nodes are iteratively removed until the removal of a node breaks the connection to the inferred targets. We use an ensemble approach where the final subnetwork is constructed by margining networks derived from each trained model, keeping only nodes and edges appearing in multiple models (see methods).

We apply this process for the case of the effects of Lestaurtinib on FOXM1, resulting in a subnetwork of the intracellular signaling network that explains this off-target effect (Figure 3E). Although strongly reduced, this network is still relatively large and difficult to interpret. This may be due to multiple plausible mechanisms being explored simultaneously as a response to limited data together with L2 regularization limitations. Alternatively, this may indicate that it is necessary to include many interactions to fully explain the off-target effect that Lestaurtinib has on FOXM1 activity, and further reduction of the network would be an oversimplification. Applying the simplest path algorithm to the network from each inferred target (in red) towards FOXM1 (in orange) we find that inhibition of CDK1 and CDK2 could lead to the direct inhibition of FOXM1 (Figure 3F). According to the model (Figure 3E), Lestaurtinib can potentially inhibit FOXM1 by inhibiting CDK1 or/and CDK2 and interacting in some uncertain manner with CDK6), meaning that the smaller subnetwork contains feasible intracellular interactions that can indeed explain the off-target effect. Indeed it has been observed that that FOXM1 can be activated by both CDK1^42^ and CDK2^42,43^, meaning their inhibition could lead to inhibition of FOXM1, as proposed by the model. Additionally, while the interaction between Lestaurtinib and CDK2 is neither present in the prior knowledge used for training nor DrugBank, it has been seen in a comprehensive kinase inhibition study that Lestaurtinib indeed inhibits CDK2 with a Kd = 20 nM^44^, which is markedly lower than the dose used in the L1000 study (10 um). We note that CDK2 was identified as an interaction in 36 out of 50 models, bordering the previously identified threshold (of 37) for identifying true direct interactions with high performance (Figure 2E-2D). CDK1 is found in 35 out of 50 models while CDK6 was found in half of the models. Taken together, this indicates that the model can be used to propose a MoA to explain the off-target effect that is biologically feasible and potentially true, which is also cell line specific, which may serve as a basis for designing therapeutic interventions or drug combinations to cancel or enhance this off-target effect.

### A case study of FOXM1 regulation by CDK2

Since the activation of FOXM1 by CDK2 and CDK1 has been experimentally demonstrated, it may serve as a useful case study for determining how well the different components in our approach recapitulate this effect. First, we inspect the inference of TF activity from gene expression data, using the DoRothEA regulon^26^ together with the VIPER algorithm^27^. For this purpose, we retrieved Affymetrix microarray data, from the Gene Expression Omnibus (GEO)^45^, generated from A375 cells treated with siRNAs against various transcription factors and signaling molecules^46^. We then inferred the activity of FOXM1 for the measured gene expression data for CDK2 and FOXM1 knockdown as well as untreated cells and control (inactive fluorescently labeled siRNAs) samples. We find that the inferred activity (z-scored) of FOXM1 when knocking down CDK2 is similar to a FOXM1 knockdown, while FOXM1 is way more inactive than in untreated cells (centered to zero as expected) in both cases (Figure 4A). Even though this published study has limited statistical power, it does indicate that our inferred activities in the L1000 recapitulate the relationship between CDK2 and FOXM1, thus corroborating the proposed off-target effect.

**Figure 4:**
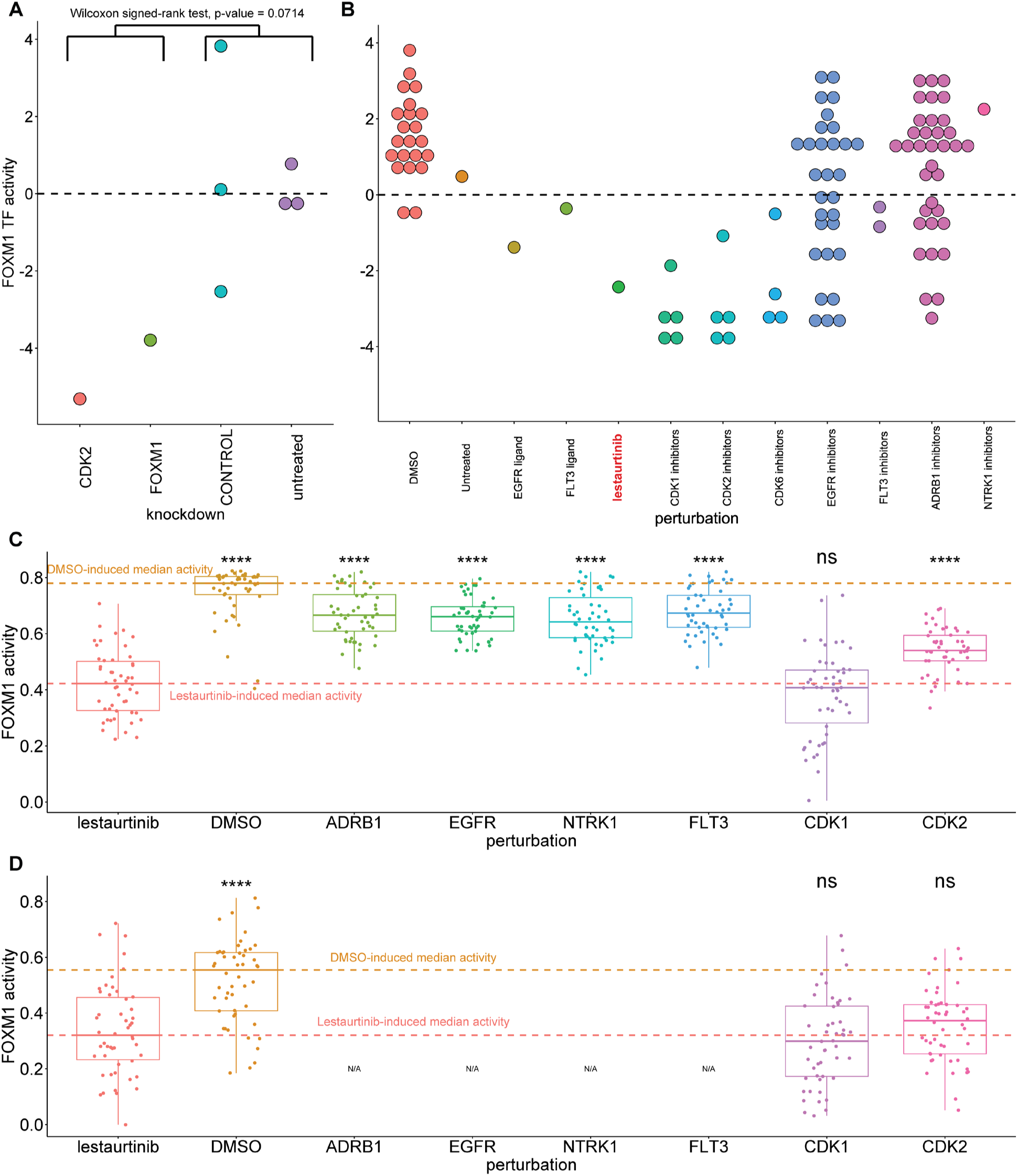
Validation of the predicted effects of Lestaurtinib on FOXM1 activity in the A375 cell line. **A)** Inferred activity after treatment with siRNA knockdowns of CDK2 and FOXM1 using public microarray data. **B)** Inferred activity after treatment with ligands and inhibitors of Lestaurtinib targets, and CDKs in the L1000 dataset. **C)** Predicted activity after an *in-silico* knockdown. **D)** Predicted FOXM1 activity using the inferred sub-network explaining the MoA of the off-target effect of Lestaurtinib. Statistical comparisons in C and D were performed relative to Lestaurtinib.

Secondly, we investigate whether the inhibition of the activity of FOXM1 by Lestaurtinib is indeed primarily achieved through inhibition of CDK2 and/or CDK1. For this purpose, we utilized data from ligand perturbations contained in the L1000 dataset but not used for training the model. From this, we inferred FOXM1 activity for ligand stimulation of the known targets of Lestaurtinib: EGFR, FLT3, and additionally for stimulation with other drugs that inhibit these targets as well as the two additional targets in DrugBank (ADRB1, NTRK1), and known inhibitors of CDK1, CDK2, and CDK6 (such as Alvocidib, AT-7519, and Kenpaullone). We made sure to select inhibitors with at most 10 targets to minimize the risk of regulation of FOXM1 through other targets. As expected, we find that the activity of FOXM1 in A375 when using Lestaurtinib is much more inhibited compared to DMSO or untreated A375 cells. We also find that it is on a similar level as for known CDK1 and/or CDK2 inhibitors (four inhibit both CDK1 and CDK2 (Figure 4B)). Meanwhile, only a few of the ADRB1 inhibitors (another Lestaurtinib target) show a similar trend, and the rest of the known targets do not inhibit FOXM1 activity at a comparable level (or not at all). This further supports the proposed MoA of Lestaurtinib inhibiting FOXM1 through the off-target effect on CDK2.

### In-Silico knockdowns using the model

Finally, we investigate if *in-*silico knockdown experiments by the model, using the proposed MoA network, recapitulate the similar effects on FOXM1 activity by Lestaurtinib, and CDK2 and CDK1 knockdowns. We induce a level of knockdown for a signaling node by assigning a large negative value as input. This way we knock down the known targets of Lestaurtinib (EGFR, NTRK1, ADRB1, FLT3) and CDK2 and CDK1, for all of the 50 trained signaling networks. Additionally, we model the signal generated by Lestaurtinib, signals masked to include either its on-target effects or off-target effects, as well as the signal from DMSO as control. We find that the activity of FOXM1 under Lestaurtinib is indeed much lower than DMSO and that seems to be mostly due to the off-target effects (Figure 4C). We note that the on-target signal induced similar activity levels as knockdowns of any of the known targets of Lestaurtinib, indicating that the model can successfully recapitulate the on-target effects of Lestaurtinib on FOXM1, which, while lower than for DMSO, do not seem to strongly inhibit FOXM1 activity (Figure 4C). Furthermore, increasing the knockdown level for these nodes does not seem to induce much stronger inhibition of FOXM1 (Supplementary Figure 11C-11D), while knocking down CDK1 and CDK2 induces strong inhibition of FOXM1, and depending on the knockdown strength (Supplementary Figure 11C-11D) induces similar inhibition as Lestaurtinib (Figure 4C) or almost completely deactivates FOXM1 (Supplementary Figure 11A-11B). Notably, if we restrict the model to only the reduced subnetwork when conducting this *in-silico* experiment, we observe a similar trend for the CDK1 and CDK2 knockdowns, suggesting that indeed the subnetwork is sufficient to explain the MoA of this off-target effect (Figure 4D). Taken together this case study serves as a proof of concept for the utilization of the models to generate *in-silico* experiments to potentially identify therapeutic interventions to cancel the off-target effect.

## Discussion

Drugs do not always function entirely through their proposed MoA^6^, which may cause adverse effects from off-targets but may also sometimes be beneficial^8,38^. Here we developed an approach for predicting the transcriptional response under drug-induced signaling, taking potential off-target effects into account. The model augments the LEMBAS framework^25^, which simulates intracellular signaling, with a trainable module for inferring drug-induced signaling, to simultaneously predict the activity of transcription factors under drug stimulation, and infer drug-target interactions that are not known. The model outperforms basic machine learning methods in predicting the transcription factor activity. It retains most of the prior knowledge of drug-target interactions but also predicts many more putative interactions, with a good balance between sensitivity and specificity. Perhaps even more importantly, we make use of integrated gradients^37^, to extract subnetworks of intracellular signaling that explain the predicted MoA of off-target effects on TF. In a case study of the drug Lestaurtinib’s off-target effects on FOXM1 activity, we demonstrated that the constructed network is biologically sensible, as we find literature support for the proposed MoA.

Understanding how the signaling effects of drugs propagate in the cell is essential for understanding how adverse effects may arise in the clinic and for designing therapy regimes that may counteract them. This is particularly important for drugs that do not function through their proposed MoA. The advent of machine learning and big data in biology holds promise for a more data-driven life science. However, machine learning models have been criticized for their lack of interpretability^47,48^ and thus many times fail to explain the underlying MoA in a biological phenomenon or were never designed to do so. Embedding prior knowledge into the structure of machine learning models can improve their interpretability ^25,49,50^. Specifically, in the case of our LEMBAS models, the whole architecture corresponds to feasible interactions in the intracellular signaling network. Combining this inherent structure with the inference of previously unknown drug-target interactions alongside a sensitivity approach to prune nodes and edges that do not contribute to its explanatory power, we were able to construct subnetworks that fully recapitulate the MoA of an off-target effect.

Despite the drastic size reduction, the subnetworks explaining the MoA of off-target effects are still far too comprehensive for immediate interpretation, and additionally, there is variation arising from the dissension between different models in the ensemble, in line with observations in the literature^51^. While it is possible that the network indeed needs to be this large to fully recapitulate the off-target effect, this may also be the result of data limitations along with the L2 regularization used to constrain the number of inferred interactions and prevent overfitting the weights of the signaling network. Multiple drug-target interactions and paths in the network might be able to explain the observed transcriptional profile. When lacking sufficient data to train a model that can fully distinguish between all feasible solutions, this can result in the model considering multiple explanations as equally important. The former would suggest that redundancy and robustness are intrinsic to cellular circuits, which implies that a reductionistic approach in biology may lead to misleading or incomplete results. The latter would indicate that while the biological process may be simple, we are currently too data-limited to confidently simplify the network further. This problem could potentially be tackled in the future either by increasing the data used for training a model, by using large transcriptomic databases such as ARCHS4^52^, or by using algorithms in the drug module that can indirectly infer interactions without the usage of L2 regularization.

In this study, the drug module, which infers drug-target interactions, is linear and relies on a pre-defined space of drugs and potential targets. It only uses knowledge about the chemical similarity of drugs, thereby ignoring potential structural similarities. However, the modular nature of our model allows for the future development of a drug module that can incorporate knowledge about targets’ structural similarity and is also non-linear. A previously proposed method, called DeepCE^16^, utilizes a graph neural network^36^ to encode the chemical structure of a drug, and an attention-based ANN^53^ to combine gene-level representations, which contain gene-gene interaction information, and drug representations in a drug-gene interaction network to ultimately predict the gene expression profile of a sample. Similarly, another approach called ChemCPA^54^ also encodes the chemical structure of the drug and non-linearly scales its dose and combines it with the drug representation. On this front, our drug module could also incorporate a non-linear encoder to represent the chemical structure of drugs and combine it with targets’ representations, by building upon ideas presented in OmegaFold^55^ and AlphaFold2^56^, in order to infer potential drug-target interactions, similar to what has been recently proposed in the ConPLex model^57^, after training models to ultimately predict the transcriptional profile of a cell.

A limitation of the present models is that they are cell line-specific and thereby do not allow an already trained model to be directly used to make predictions for other cell lines. Moreover, when inferring drug-target interactions solely based on one cell line-specific model, we might miss potential general drug-target interactions, as a target molecule may not be expressed in this particular cellular model, thus resulting in more false negatives. Finding a basal representation for each cell line, like the sequencing profile of cell lines from the CCLE database^58^, and using that as input to the model, could be used to train a unified model that also incorporates cell lines as input in a non-predefined cell line space. Generally, contextualizing a unified model or transferring predictions and MoA representation from one cellular model to another would be important for the utility of the model.

Our framework introduces a way to conduct *in-silico* experiments of drug perturbations while simultaneously being able to explain the MoA of a drug. As such, future use may be for designing drug combination therapies while exploring and studying their synergistic or competitive effects, identifying ways to counter drugs’ off-target effects, and designing better therapeutic regimes with higher clinical efficacy.

## Materials and Methods

### Retrieving prior knowledge networks of drug-target interactions

Drug-target interactions for training our models were retrieved from Broad’s Institute Repurposing Hub^33^. The network was subset to drugs with corresponding perturbations in the L1000 dataset^14^. Drugs were mapped with their respective targets by multiple identifiers for the drugs, namely: 1) the drugs’ SMILEs, 2) the International Chemical Identifier (InChIKey), 3) the PubChem Compound Identifier (pubchem_cid), 4) the Broad’s Institute internal identifier (pert_id), and 5) the drugs’ common names. The targets for DMSO were manually curated from DrugBank^13^. For the evaluation of the model’s ability to retrieve drug-target interactions, we retrieved additional interactions from DrugBank^13^, using drugs’ common names.

### Pre-processing of *in-vitro* transcriptomics in the L1000 dataset

Transcriptomic signatures of drug perturbations were retrieved from the L1000 dataset^14^ (accessed via GEO with accession number: GSE92742). For inferring TF activity, we utilized gene expression data of 978 landmark genes, measured with the L1000 assay, and additionally, 9,196 imputed genes that were labeled as well-inferred by the L1000 study^14^. The data were retrieved at Level 3, one of the processing steps in the pipeline of the L1000 dataset, containing normalized gene expression data. We considered only *exemplar* signatures, which, according to the L1000 definition, are the signatures with highest the transcriptional activity score (TAS) in the case of multi-signature perturbagens, i.e. technical duplicates. Briefly, the TAS metric inherent to the L1000 dataset quantifies signal strength and reproducibility, and definitions and further information is available in the CLUE platform^59^ glossary. Additionally, we keep only drugs with at least one known target in the prior knowledge signaling network (see methods section for constructing the prior knowledge signaling network). After inferring TF activity and further filtering data to keep only high-quality TF activity data, we keep perturbations with at least 400 unique drugs per cell line (the number of conditions previously found to achieve high performance when training a LEMBAS signaling model^25^). This filtering results in 9 cancer cell lines and for evaluation purposes (see evaluation method section), resulting in a drug space of 233 unique drugs. The log-scaled dose is used as input to train models (*dose_scaled_* = log_10_(*dose* + 1)). For the *in-silico* validation case study of Lestaurtinib, we utilized the level 5 z-score transformed data were replicates are already aggregated, and specifically for shRNA, ligand, and control (DMSO-treated and untreated cells) data we kept aggregated signatures derived from at least 3 technical replicates.

### Pre-processing of *in-vitro* Affymetrix microarray data

For the publicly available siRNA experiments^46^, we retrieved Affymetrix microarray data from the Gene Expression Omnibus (GEO)^45^, under the GSE31534 ascension number. The raw microarray gene expression data were normalized using the Robust Multichip Averaging (RMA) algorithm^60^ included in the *affy* R package^61^. The normalized expression values were used to infer TF activity (see below).

### Inference and pre-processing of transcription factor activity data

The activity of transcription factors (TFs) was inferred from transcriptomics data using the VIPER algorithm^27^ coupled with the Dorothea regulon^26^. The VIPER algorithm calculates the enrichment of known regulons (TFs), which act as proxies of TF activity. The activity of a TF is calculated based on the expression of downstream genes known to be regulated by this specific TF, utilizing a known transcription regulatory network. The Dorothea regulon contains known regulatory interactions and thus can be used to build a regulatory network. Here we kept only high-confidence interactions (confidence levels A and B).

After inferring the TF activity of the pre-processed transcriptomic data in the L1000 dataset, we filtered TFs with high variance across technical replicates, to ensure we kept only high-quality estimations of TF activity, and then we filtered technical replicates that were not correlated enough with the other replicate signatures. To filter TFs, we first build a null distribution of TF activity variance by permuting 100 times the rows (samples) of the activity matrix, labeling this way random profiles as technical replicates, and then calculating the variance of the activity of each TF across each group of replicates. This way a null distribution of TF activity variances is built for each TF. The actual distribution of TF activity variances across replicates is compared with the null distribution, using a one-tailed Kolmogorov–Smirnov statistical test, to test whether the actual variance across replicates is less than the random variance. If the p-value is greater than 0.05 the tested TF is removed and will not be utilized in downstream analysis. To filter replicates, we build a null distribution of random correlations between TF activity profiles, by randomly sampling 1000 times an equal number of signatures as the number of replicates, calculating the Pearson’s correlation between each pair and taking the mean correlation as a proxy of how similar the replicates are within a sample. We repeat this for every possible number of replicates within a sample. Then we calculate the correlation between each actual technical replicate with all others in a sample and count how many random correlations are equal to or higher than the mean correlation of the technical replicates, to calculate the probability of observing a given correlation due to change. If the p-value is more than 0.05 we remove the sample and all of its replicates. Finally, we merge replicate signatures by using the median of their TF activity profiles. In case there is only one replicate, we keep the sample as it is.

### Reconstructing a prior knowledge of signaling network

We reconstructed a prior knowledge intracellular signaling network (PKN), to constrain our ANN signaling model, from protein-protein interactions retrieved from the OmniPath database^20^. Only human interactions from the OmniPath core set were included and further restricted to interactions originating either from the KEGG^62^, InnateDB^63^, or SIGNOR^21^ resources. First of all, we remove TFs and drug targets not included in the core prior knowledge network. Then we trim the PKN by removing nodes and edges from the network if for some nodes there was no path from any drug to any TF. Additionally, nodes were removed if they had only a single source and target that both were the same node. Finally, we removed TFs and drug-target interactions if a target or TF is not in the final trimmed PKN. Drugs that remained with no target in the constructed prior knowledge are removed from our data used to train and validate the model.

### Model architecture

The model consists of two interconnected modules. First, a drug module that takes as input the concertation of a drug, in a pre-defined drug-target space, infers drug signaling. This utilizes the known drug-target interactions (*W*_*DT*_) and the pre-calculated chemical similarity (denoted as *W*_*sim*_ with [d x d] dimensions, where d is the number of drugs available), using the tanimoto similarity of drugs ECFP4 fingerprints^34^, between drugs in the drug space. Ultimately the drug signaling (*S* with [n x t] dimensions, where n is the number of conditions and t is the number of available targets) which is the output of the drug module is given by: *S* = *bn* (*X* ∗ (*W*_*sim*_ ⊙ *W*_*drug*_)) ∗ *W*_*DT*_. Specifically, the input concentration matrix (*X*) of available drugs is first multiplied by the element-wise product between the pre-calculated chemical similarity and a trainable weight matrix (*W*_*drug*_), acting as a trainable scaler of chemical similarity, and thus controlling to which extent chemical similarity should contribute to the models’ predictions. The result of this operation is passed through a batch normalization layer^64^ with a momentum of 0.6, and, during training only, a dropout layer^65^, with a drop-out rate of 0.1. Finally, it is multiplied with a sparse trainable weight matrix (*W*_*DT*_) containing known drug-target interactions (dimensions [d x t]). The drug signaling (*S*) generated by the drug module, which represents the signal created by the drugs in a pre-defined drug-target space, is used as the input to the second module.

The second module is the LEMBAS framework^25^ which contains a recurrent ANN model of intracellular signaling, where the connections are based on prior knowledge of the intracellular signaling network. In LEMBAS the signaling state of each node is calculated using the signaling state of the interacting node in the previous time step, by multiplying it with a trainable connectivity matrix and adding a trainable bias, all passed through a non-linear Michaelis–Menten-like (MML) activation function, as proposed in the LEMBAS manuscript^25^. Drug signaling (*S*) is first projected on the signaling nodes’ space and it is used as input in the LEMBAS network. The state vector, describing the signaling state of each node, is initialized as all 1e-3, except for TF nodes which are initialized as 0.5, and iterated for a maximum of 120 steps, after which it is assumed that a steady state has been reached. Finally, the TF activity is predicted by projecting from the signaling state of the network at the steady state.

### Training of the model

A cell line-specific model is trained for 5000 epochs to ultimately predict the activity of 101 TFs, given the concentration of a drug, in a pre-defined drug-target space of 233 drugs and 259 potential targets. The term describing the main task of the model during training (*fitLoss*) is given by the Mean Squared Error (MSE) across TFs, averaged across a batch (batch size=25) of data points used to update the weights of the model during a learning cycle. There are auxiliary terms in the training loss of the model, to constrain different parts of it, and we incorporated them from the GitHub repository (https://github.com/Lauffenburger-Lab/LEMBAS) of the LEMBAS framework^25^. More details are included in Supplementary Note 1, but we briefly list the regularization terms in this method section. The *signConstraint* constrains the sign of the weights in the intracellular network to be the same as the sign of known protein-protein interactions. The weights and biases of the intracellular signaling network are L2-regularized with the *NetWeightLoss* and *biasLoss* terms respectively. The trainable weights used to project from the signaling state to TF activity were also L2-regularized in the *projectionLoss* term. Moreover, specific statistical properties for the activities of the signaling nodes are enforced with the *stateLoss* term. Finally, following the implementation proposed in the LEMBAS framework^25^, to ensure that the model achieves convergence by reaching a steady state we aim to constrain the absolute value of the largest eigenvalue of the transition matrix, i.e., the spectral radius (ρ), to be less than 1 in the *spectralRadiusLoss* term (more details in Supplementary Note 1).

For the drug module, we implement two additional terms. First, we treat the drug-target interaction matrix as a small network and we regularize the weights similar to what we have done in the signaling network: 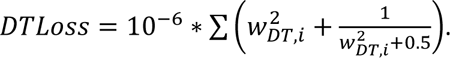. Secondly, we implement a regularization term (*DTregularization*) using the trainable *W*_*drug*_ matrix (described in the previous section), to control how many new interactions should be inferred and, thus how much should the model be allowed to deviate from prior knowledge by considering chemical similarity (more details in the following corresponding section). The final formula describing the total training loss, which is minimized by updating the model’s parameters using the Adam optimizer^66^ with a learning rate ranging from 10^-8^ to 2*10^-3^ is:

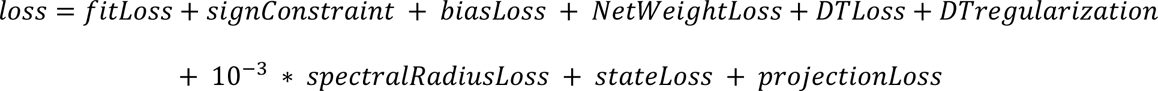

### Evaluation of the model

To evaluate the generalization of the drug module and the LEMBAS part of the framework to unseen conditions, we implement a validation procedure where we train a whole model in the cell line with the most conditions available (VCAP), we freeze the weights of the drug module, and re-train only the signaling network part, in every one of the other 8 remaining cell lines, by using only 80% of the available drugs, while we make sure that the 20% hidden are drugs dissimilar from the ones used in training (regarding their chemical structure). If the drug module is not general enough the signaling network may change a lot and fail to generalize in dissimilar cases.

### Regularization of the inference of drug-target interactions

To constrain the number of inferred drug-target interactions we regularize the weights of the previously described *W*_*drug*_ matrix, containing trainable weights to scale the similarity between the available drugs in our data, such as that *W*_*drug*_ is closer to the identity matrix (*I*). Thus, the regularization term used in the loss function is formed as:

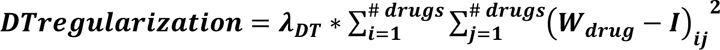

Where *λ*_*DT*_ is a free user-defined parameter, quantifying the strength of regularization. In this study, we performed an analysis, to study the effect of regularizing the drug-target interactions inference, with testing values from zero to infinity, where infinity, means we train a model using only the sparse trainable weight matrix (*W*_*DT*_) containing known drug-target interactions. Since, the operation between *W*_*drug*_ and *W*_*sim*_, containing pre-calculated chemical similarity, is that of element-wise multiplication, if *W*_*drug*_ = *I*, then *W*_*sim*_ ⊙ *W*_*drug*_ = *I*, meaning that the output of the drug module degenerates to: *S* = *X* ∗ *W*_*DT*_, meaning using only prior knowledge of drug-target interactions, which theoretically would be achieved with infinite regularization (*λ*_*DT*_ → ∞).

### The drug-target interaction inference algorithm

To infer drug-target interactions using the drug module, how much a drug affects a potential target is quantified by using integrated gradients^37^ from the Captum library^67^: 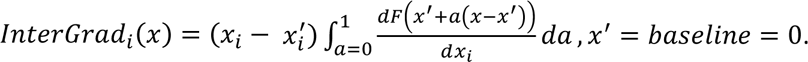 To identify a cut-off for identifying significantly large scores we utilize an error-based approach where we calculate the Mean Absolute Error (MAE) of the model across all TFs, after removing drug-target interactions, and thus drug input signal in LEMBAS, for increasingly higher absolute gradient score. We select as a cut-off the score that induces a 25% (or larger) increase in the model’s MAE. Drug-target interactions with a smaller score than the cut-off are considered insignificant, and thus are disregarded. Finally, we utilize the ensemble of models to derive a frequency score for each interaction appearing in multiple models and further filter the inferred drug-target interactions.

### Node and edge importance in affecting a specific TF

To quantify the importance of a node or an edge in regulating the activity of a TF of interest we utilize a customized integrated gradient approach. First, we generate for each model the input signal from the drug module, and then we pass through the signaling module fractions of this signal’s strength, ranging from 0 to 1. We denote this input matrix *X*_*in*_. The sum of the TF activity across all these artificial conditions is used as an objective function (*L*_*obj*_) for which the gradients for the weights (*dw*) and biases (*db*) of the signaling module, are calculated using back-propagation. The node importance was calculated as: *score*_*b*_= |*db*| ∗ |*range*|, where *range* is the range of the node activity, for different signals’ strength. The edge importance was calculated as: *score*_*w*_= |*dw*| ∗ |*weight*|, where *weight* is the weight of an edge in the model. More details about the implementation and calculation of each term can be found in Supplementary Note 2.

### Identifying samples with high off-target effect

We remove the off-target signal and only the signal on the known targets is used to predict TF activities. The difference between the original predictions of the model and the ones where off-targets are masked out (Δactivity) quantifies the magnitude of the off-target effects on the TF of interest. The calculated Δactivity is derived from the mean TF activity prediction from an ensemble of 50 trained models. Samples where a TF has an activity≥0.75 or activity≤0.25, |Δ*activity*| ≥ 0.2, average Pearson’s r (between training and validation) of at least 0.5, and average validation Pearson’s greater than 0.4, are considered trustworthy predictions with a large off-target effect on a specific TF. For the second step, we infer drug-target interactions for each model as previously discussed.

### Algorithm for subsetting the network to the mechanism of action

To subset the signaling network for explaining the MoA of off-target effects edges are removed from the whole signaling network based on their importance in regulating the activity of a TF of interest. Nodes and edges are removed iteratively based on their importance (see previous section) until further removal results in the removal of all target nodes or until there is no path from the drug’s target to the TF of interest. First, we remove nodes and get rid of disconnected parts of the network, nodes that the drug cannot access through any path, and paths whose end is not the drug’s target or the TF. Then we remove edges and repeat the aforementioned network cleaning. The drug’s targets with no path to the TF are removed. Finally, we keep inferred targets that appear in at least 50% of the models, if possible, otherwise, we use a cut-off that results in at least one inferred target in that subnetwork. We do the same for edges, but if there is not a single edge that can be removed based on some frequency threshold, without maintaining the connection between some target and the TF, we use a threshold of 50% and then we start including gradually more edges to connect some target with the TF, and we keep the edges of the path with the highest sum of frequency scores (regarding the frequency in appearing in multiple models). In every part of this final trimming process, we also perform basic cleaning of the network by removing undruggable nodes, nodes that cannot affect the TF via some path, and disconnected parts of the network.

### In-silico knockouts

To induce *in-silico* knockouts of signaling nodes, and to validate different MoAs and off-target effects, we assign a largely negative value in the input signal, that is used as input in the trained LEMBAS part of the model, to the node we wish to induce a knockout. Then this signal is propagated in the network and the model iterates for at least 120 steps, or until convergence.

### Lethality predictions using drug-target interactions

Random Forest models (RF) are trained to predict the lethality of drug perturbations on cancer cell lines, by using as input drug-target interactions and the cancer cell line on which the drug was tested, as a one-hot encoded vector. Lethality data of drugs tested on different 9 cell lines in our study from the NCI60 drug screen^39^ were accessed via the PharmacoDB database^68,69^. Separate RF models were trained and tested using only the prior knowledge of drug-target interactions used in the drug module of our framework and then using the inferred interaction. Data from the A549 cancer cell were used for validation and the rest were used to train the RF models.

### Statistics and reproducibility

For the evaluation of performance in retrieving drug-target interactions (in Figure 2), metrics and the p-values were calculated, inside the *caret* R package^70^, with a binomial one-tailed test comparing the proportions of accuracy and NIR^71^. Statistical comparisons of models’ performance in terms of Pearson’s correlation are conducted using a two-sided unpaired Wilcoxon test. Non-parametric Kolmogorov– Smirnov tests are used to compare whole distributions (see the corresponding sections when they are used for more details).

### Hardware and software specifications

All models were expressed in and trained using the PyTorch framework^72^ (versions 1.10.2 & 1.12) in Python (version 3.6.13 & 3.8.8). Generally, simple simulations using one model were performed on a Dell XPS 17 laptop with an Intel i9-11900h @4.9 GHz with 8 cores (16 logic processors) and 32 GB RAM. For convenience, ensemble training of multiple models, random models training, and cross-validation was carried out on a single-threaded computer cluster (Intel Xeon CPU @ 2.60 GHz) that allowed job scheduling (using Slurm) with 16 parallel jobs. Pre-processing and statistical analysis of the results were done in the R programming language (version 4.1.2). Visualization of results was done mainly using ggplot2^73^. More information about the versions of each library used can be found in the GitHub provided in the Data and Code Availability section.

## Supporting information

DTLEMBAS_supplementary

## Data and Code availability

The study did not produce new experimental data. All analyzed data that were used to train our models and produce all tables and figures, as well as all the code to generate those data, figures, and tables are available at the following GitHub repository: https://github.com/Lauffenburger-Lab/DrugsANNSignaling

## Acknowledgments

The authors would like to thank Brian Joughin, Diana Gong, Christine Wiggins, Krista Pullen, Anisha Datta, Jose Cadavid, Michal Caspi Tal, Erin Tevonian, and Andy Lopez for their valuable input on this work. We acknowledge funding from the Swedish Research Council, grant no. 2019-06349 and the SciLifeLab & Wallenberg Data Driven Life Science Program grant no. KAW 2020.0239 (AN). We also acknowledge funding from US ARO cooperative agreement W911NF-19-2-0026 for the Institute for Collaborative Biotechnologies (DAL) and NIH contract NIH contract #75N93019C00071 (DAL).

## References

1. Plati, J., Bucur, O. & Khosravi-Far, R. Dysregulation of apoptotic signaling in cancer: Molecular mechanisms and therapeutic opportunities. Journal of Cellular Biochemistry 104, 1124–1149 (2008).

2. Wu, W. K. K. et al. Dysregulation of cellular signaling in gastric cancer. Cancer Letters 295, 144–153 (2010).

3. García-Velázquez, L. & Arias, C. The emerging role of Wnt signaling dysregulation in the understanding and modification of age-associated diseases. Ageing Research Reviews 37, 135–145 (2017).

4. Popugaeva, E., Pchitskaya, E. & Bezprozvanny, I. Dysregulation of Intracellular Calcium Signaling in Alzheimer’s Disease. Antioxidants & Redox Signaling 29, 1176–1188 (2018).

5. Adjei, A. A. Blocking Oncogenic Ras Signaling for Cancer Therapy. JNCI: Journal of the National Cancer Institute 93, 1062–1074 (2001).

6. Lin, A. et al. Off-target toxicity is a common mechanism of action of cancer drugs undergoing clinical trials. Science Translational Medicine 11, eaaw8412 (2019).

7. Bai, J. P. F. & Abernethy, D. R. Systems Pharmacology to Predict Drug Toxicity: Integration Across Levels of Biological Organization. Annual Review of Pharmacology and Toxicology 53, 451–473 (2013).

8. Hopkins, A. L. Network pharmacology: the next paradigm in drug discovery. Nat Chem Biol 4, 682–690 (2008).

9. Fotis, C., Antoranz, A., Hatziavramidis, D., Sakellaropoulos, T. & Alexopoulos, L. G. Network-based technologies for early drug discovery. Drug Discovery Today 23, 626–635 (2018).

10. Verbist, B. et al. Using transcriptomics to guide lead optimization in drug discovery projects: Lessons learned from the QSTAR project. Drug Discovery Today 20, 505–513 (2015).

11. Yang, X. et al. High-Throughput Transcriptome Profiling in Drug and Biomarker Discovery. Frontiers in Genetics 11, (2020).

12. Schenone, M., Dančík, V., Wagner, B. K. & Clemons, P. A. Target identification and mechanism of action in chemical biology and drug discovery. Nat Chem Biol 9, 232–240 (2013).

13. Wishart, D. S. et al. DrugBank: a comprehensive resource for in silico drug discovery and exploration. Nucleic Acids Research 34, D668–D672 (2006).

14. Subramanian, A. et al. A Next Generation Connectivity Map: L1000 Platform and the First 1,000,000 Profiles. Cell 171, 1437–1452.e17 (2017).

15. Douglass, E. F. et al. A community challenge for a pancancer drug mechanism of action inference from perturbational profile data. Cell Reports Medicine 3, 100492 (2022).

16. Pham, T.-H., Qiu, Y., Zeng, J., Xie, L. & Zhang, P. A deep learning framework for high-throughput mechanism-driven phenotype compound screening and its application to COVID-19 drug repurposing. Nat Mach Intell 3, 247–257 (2021).

17. Pritchard, J. R., Bruno, P. M., Hemann, M. T. & Lauffenburger, D. A. Predicting cancer drug mechanisms of action using molecular network signatures. Mol. BioSyst. 9, 1604–1619 (2013).

18. Hyduke, D. R. & Palsson, B. Ø. Towards genome-scale signalling-network reconstructions. Nat Rev Genet 11, 297–307 (2010).

19. Münzner, U., Lubitz, T., Klipp, E. & Krantz, M. Toward Genome-Scale Models of Signal Transduction Networks. in Systems Biology 215–242 (John Wiley & Sons, Ltd, 2017). doi:10.1002/9783527696130.ch8.

20. Türei, D., Korcsmáros, T. & Saez-Rodriguez, J. OmniPath: guidelines and gateway for literature-curated signaling pathway resources. Nat Methods 13, 966–967 (2016).

21. Lo Surdo, P., et al. SIGNOR 3.0, the SIGnaling network open resource 3.0: 2022 update. Nucleic Acids Research 51, D631–D637 (2023).

22. Saez-Rodriguez, J. et al. Discrete logic modelling as a means to link protein signalling networks with functional analysis of mammalian signal transduction. Molecular Systems Biology 5, 331 (2009).

23. Fröhlich, F. et al. Efficient Parameter Estimation Enables the Prediction of Drug Response Using a Mechanistic Pan-Cancer Pathway Model. Cell Systems 7, 567–579.e6 (2018).

24. Fortelny, N. & Bock, C. Knowledge-primed neural networks enable biologically interpretable deep learning on single-cell sequencing data. Genome Biology 21, 190 (2020).

25. Nilsson, A., Peters, J. M., Bryson, B. & Lauffenburger, D. A. Artificial neural networks enable genome-scale simulations of intracellular signaling. 2021.09.24.461703 https://www.biorxiv.org/content/10.1101/2021.09.24.461703v1 (2021) doi:10.1101/2021.09.24.461703.

26. Garcia-Alonso, L., Holland, C. H., Ibrahim, M. M., Turei, D. & Saez-Rodriguez, J. Benchmark and integration of resources for the estimation of human transcription factor activities. Genome Res. 29, 1363–1375 (2019).

27. Alvarez, M. J. et al. Functional characterization of somatic mutations in cancer using network-based inference of protein activity. Nat Genet 48, 838–847 (2016).

28. Bonner, S. et al. A Review of Biomedical Datasets Relating to Drug Discovery: A Knowledge Graph Perspective. Briefings in Bioinformatics 23, bbac404 (2022).

29. Gupta, R. et al. Artificial intelligence to deep learning: machine intelligence approach for drug discovery. Mol Divers 25, 1315–1360 (2021).

30. Drug discovery with explainable artificial intelligence | Nature Machine Intelligence. https://www.nature.com/articles/s42256-020-00236-4#Sec3.

31. Öztürk, H., Özgür, A. & Ozkirimli, E. DeepDTA: deep drug–target binding affinity prediction. Bioinformatics 34, i821–i829 (2018).

32. Nguyen, T. et al. GraphDTA: predicting drug–target binding affinity with graph neural networks. Bioinformatics 37, 1140–1147 (2021).

33. Corsello, S. M. et al. The Drug Repurposing Hub: a next-generation drug library and information resource. Nat Med 23, 405–408 (2017).

34. Rogers, D. & Hahn, M. Extended-Connectivity Fingerprints. J. Chem. Inf. Model. 50, 742–754 (2010).

35. Wallach, I. & Heifets, A. Most Ligand-Based Classification Benchmarks Reward Memorization Rather than Generalization. J. Chem. Inf. Model. 58, 916–932 (2018).

36. Duvenaud, D. et al. Convolutional Networks on Graphs for Learning Molecular Fingerprints. arXiv:1509.09292 [cs, stat] (2015).

37. Sundararajan, M., Taly, A. & Yan, Q. Axiomatic Attribution for Deep Networks. in Proceedings of the 34th International Conference on Machine Learning 3319–3328 (PMLR, 2017).

38. Gujral, T. S., Peshkin, L. & Kirschner, M. W. Exploiting polypharmacology for drug target deconvolution. Proceedings of the National Academy of Sciences 111, 5048–5053 (2014).

39. Shoemaker, R. H. The NCI60 human tumour cell line anticancer drug screen. Nat Rev Cancer 6, 813–823 (2006).

40. Siddarth, V. & Gujral, T. Non-Linear Deep Neural Network for Rapid and Accurate Prediction of Phenotypic Responses to Kinase Inhibitors. SSRN Scholarly Paper at 10.2139/ssrn.3541363 (2020).

41. Liao, G.-B. et al. Regulation of the master regulator FOXM1 in cancer. Cell Communication and Signaling 16, 57 (2018).

42. Wierstra, I. & Alves, J. FOXM1c is activated by cyclin E/Cdk2, cyclin A/Cdk2, and cyclin A/Cdk1, but repressed by GSK-3α. Biochemical and Biophysical Research Communications 348, 99–108 (2006).

43. Lüscher-Firzlaff, J. M., Lilischkis, R. & Lüscher, B. Regulation of the transcription factor FOXM1c by Cyclin E/CDK2. FEBS Letters 580, 1716–1722 (2006).

44. Davis, M. I. et al. Comprehensive analysis of kinase inhibitor selectivity. Nat Biotechnol 29, 1046– 1051 (2011).

45. Edgar, R., Domrachev, M. & Lash, A. E. Gene Expression Omnibus: NCBI gene expression and hybridization array data repository. Nucleic Acids Research 30, 207–210 (2002).

46. Wang, L. et al. Cell Cycle Gene Networks Are Associated with Melanoma Prognosis. PLOS ONE 7, e34247 (2012).

47. Ching, T. et al. Opportunities and obstacles for deep learning in biology and medicine. Journal of The Royal Society Interface 15, 20170387 (2018).

48. Wysocka, M., Wysocki, O., Zufferey, M., Landers, D. & Freitas, A. A systematic review of biologically-informed deep learning models for cancer: fundamental trends for encoding and interpreting oncology data. BMC Bioinformatics 24, 198 (2023).

49. Lotfollahi, M. et al. Biologically informed deep learning to query gene programs in single-cell atlases. Nat Cell Biol 25, 337–350 (2023).

50. Integrating knowledge and omics to decipher mechanisms via large-scale models of signaling networks | Molecular Systems Biology. https://www.embopress.org/doi/full/10.15252/msb.202211036.

51. Esser-Skala, W. & Fortelny, N. Reliable interpretability of biology-inspired deep neural networks. 2023.07.17.549297 Preprint at 10.1101/2023.07.17.549297 (2023).

52. Lachmann, A. et al. Massive mining of publicly available RNA-seq data from human and mouse. Nat Commun 9, 1366 (2018).

53. Vaswani, A. et al. Attention is all you need. in Advances in neural information processing systems 5998–6008 (2017).

54. Hetzel, L. et al. Predicting Cellular Responses to Novel Drug Perturbations at a Single-Cell Resolution. Advances in Neural Information Processing Systems 35, 26711–26722 (2022).

55. High-resolution de novo structure prediction from primary sequence | bioRxiv. https://www.biorxiv.org/content/10.1101/2022.07.21.500999v1.abstract.

56. Jumper, J. et al. Highly accurate protein structure prediction with AlphaFold. Nature 596, 583– 589 (2021).

57. Singh, R., Sledzieski, S., Bryson, B., Cowen, L. & Berger, B. Contrastive learning in protein language space predicts interactions between drugs and protein targets. Proceedings of the National Academy of Sciences 120, e2220778120 (2023).

58. Ghandi, M. et al. Next-generation characterization of the Cancer Cell Line Encyclopedia. Nature 569, 503–508 (2019).

59. [clue.io]. https://clue.io/.

60. Irizarry, R. A. et al. Summaries of Affymetrix GeneChip probe level data. Nucleic Acids Research 31, e15 (2003).

61. Gautier, L., Cope, L., Bolstad, B. M. & Irizarry, R. A. affy—analysis of Affymetrix GeneChip data at the probe level. Bioinformatics 20, 307–315 (2004).

62. Kanehisa, M. & Goto, S. KEGG: Kyoto Encyclopedia of Genes and Genomes. Nucleic Acids Research 28, 27–30 (2000).

63. Lynn, D. J. et al. InnateDB: facilitating systems-level analyses of the mammalian innate immune response. Molecular Systems Biology 4, 218 (2008).

64. Ioffe, S. & Szegedy, C. Batch Normalization: Accelerating Deep Network Training by Reducing Internal Covariate Shift. in Proceedings of the 32nd International Conference on Machine Learning 448–456 (PMLR, 2015).

65. Srivastava, N., Hinton, G., Krizhevsky, A., Sutskever, I. & Salakhutdinov, R. Dropout: A Simple Way to Prevent Neural Networks from Overfitting. Journal of Machine Learning Research 15, 1929–1958 (2014).

66. Kingma, D. P. & Ba, J. Adam: A Method for Stochastic Optimization. arXiv.org https://arxiv.org/abs/1412.6980v9 (2014).

67. Kokhlikyan, N., et al. Captum: A unified and generic model interpretability library for PyTorch. Preprint at 10.48550/arXiv.2009.07896 (2020).

68. Smirnov, P. et al. PharmacoDB: an integrative database for mining in vitro anticancer drug screening studies. Nucleic Acids Research 46, D994–D1002 (2018).

69. Smirnov, P. et al. PharmacoGx: an R package for analysis of large pharmacogenomic datasets. Bioinformatics 32, 1244–1246 (2016).

70. Kuhn, M. Building Predictive Models in R Using the caret Package. Journal of Statistical Software 28, 1–26 (2008).

71. Bicego, M. & Mensi, A. Null/No Information Rate (NIR): a statistical test to assess if a classification accuracy is significant for a given problem. Preprint at 10.48550/arXiv.2306.06140 (2023).

72. Paszke, A. et al. PyTorch: An Imperative Style, High-Performance Deep Learning Library. In Advances in Neural Information Processing Systems vol. 32 (Curran Associates, Inc., 2019).

73. Villanueva, R. A. M. & Chen, Z. J. ggplot2: Elegant Graphics for Data Analysis (2nd ed.). Measurement: Interdisciplinary Research and Perspectives 17, 160–167 (2019).

