## Supplementary material for "Inference of drug off-target effects on cellular signaling using Interactome-Based Deep Learning": DTLEMBAS_supplementary

### Supplementary Figures

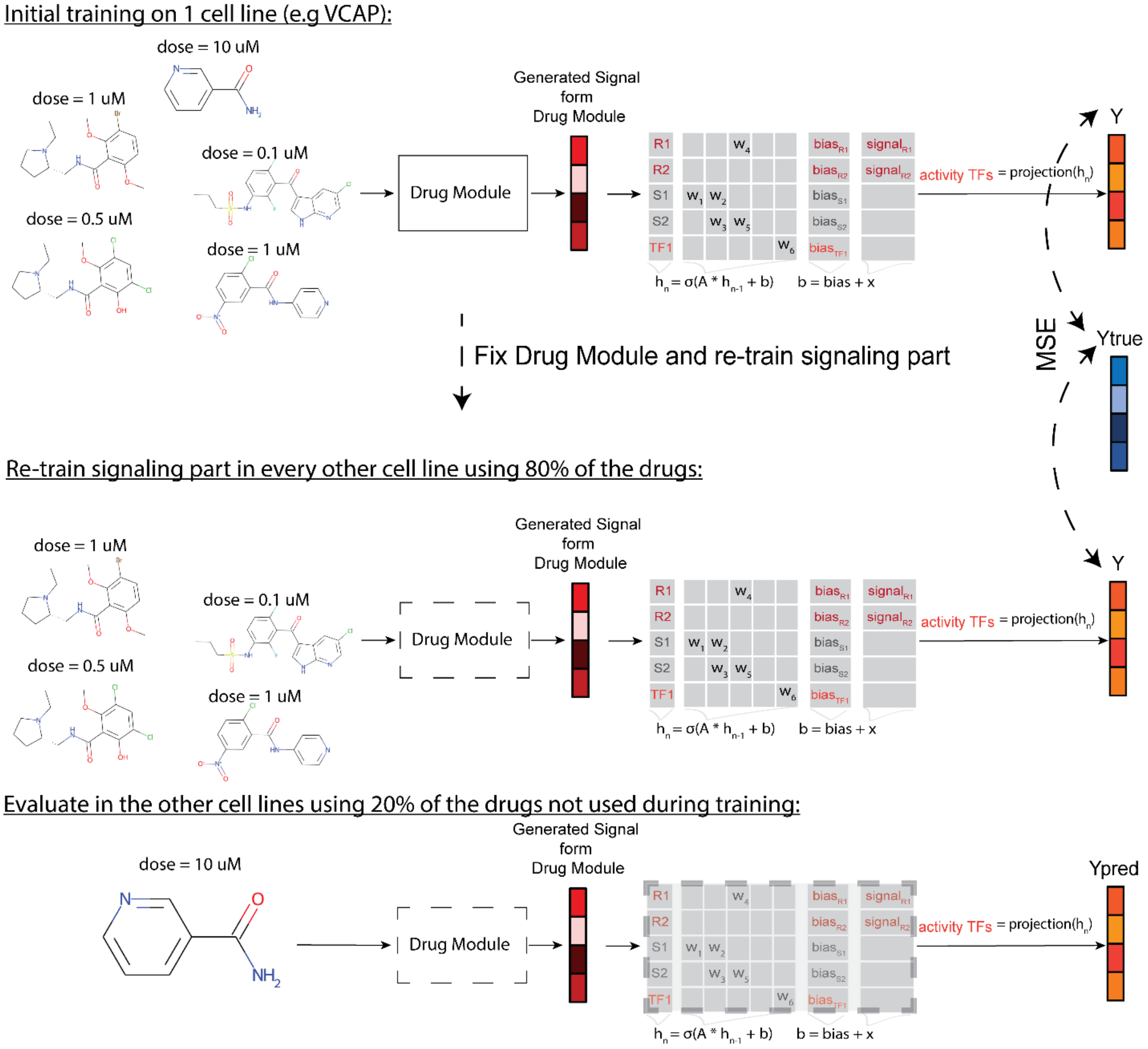


**Supplementary Figure 1: Training and validation procedure**

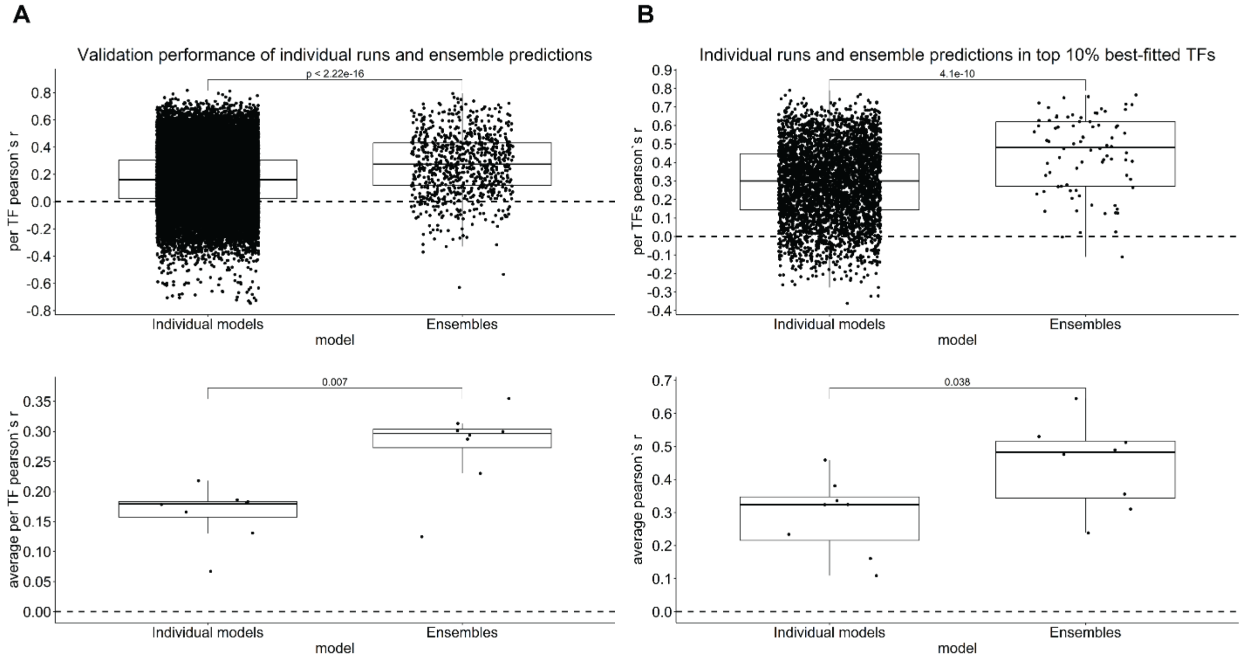


**<u>Supplementary Figure 2: Performance comparison of the ensemble approach and individual models, using the VCAP cell line for initial training</u> A)** Performance comparison of the ensemble approach and the individual models, using Pearson's r between predicted and actual TF activity **B)** Performance comparison of the ensemble approach and the individual models, by looking only at the top 10% well-fitted TFs during training.

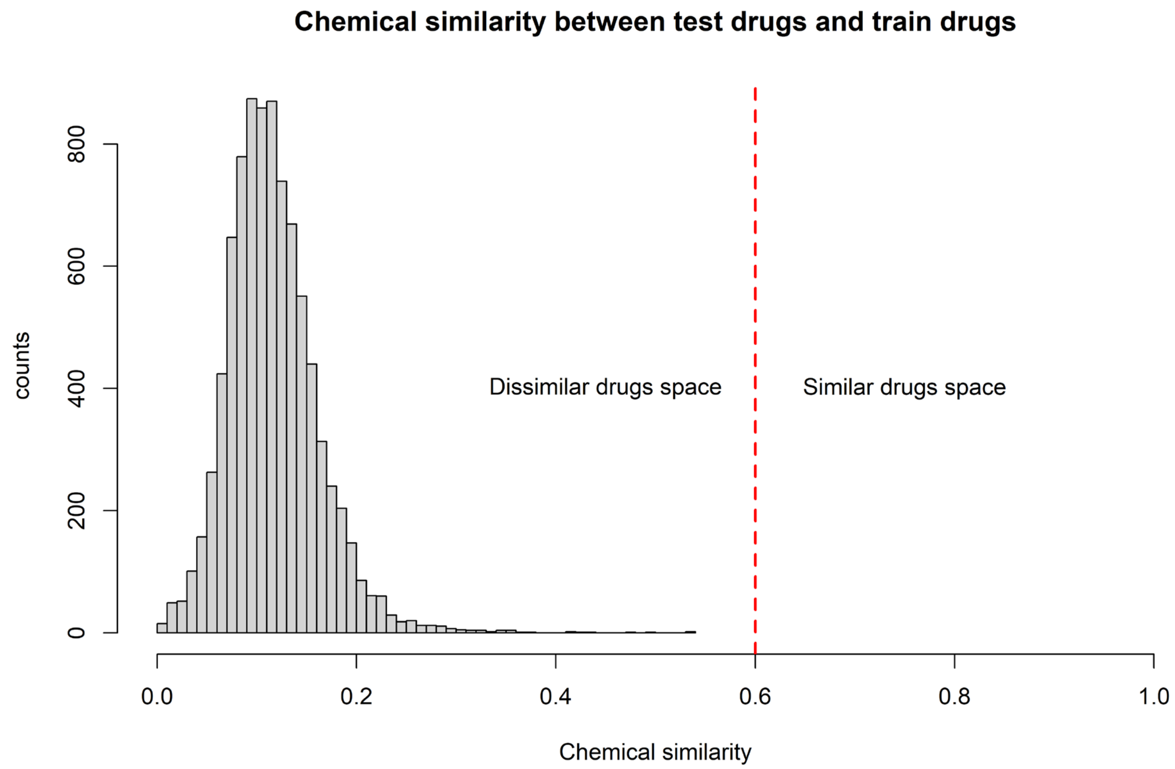


**<u>Supplementary Figure 3: Chemical similarity of validation and train drugs.</u>** Distribution of the Tanimoto similarity between ECFP4 fingerprints (chemical similarity) of any validation drug with any other drug used in training the models.

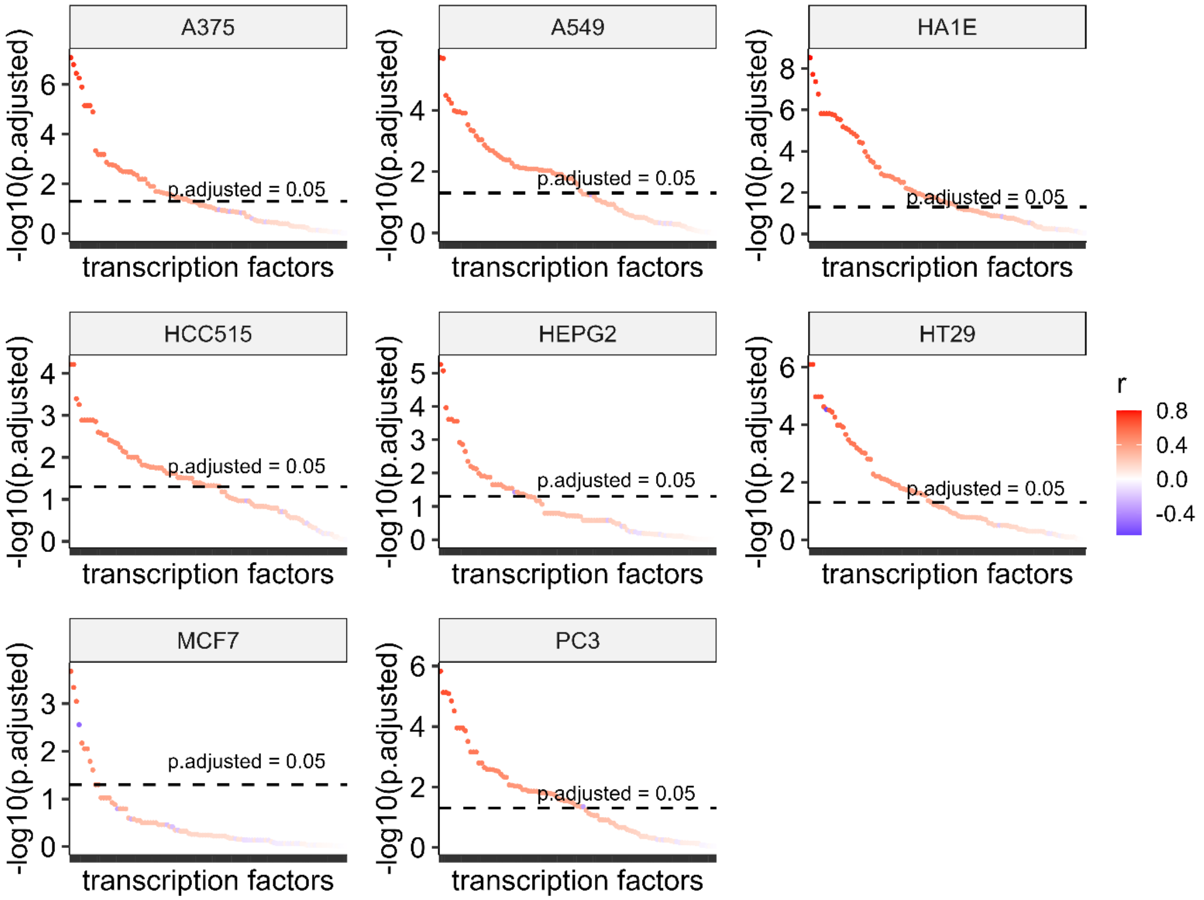


**<u>Supplementary Figure 4: Adjusted p-values of the performance of individual TFs in the ensemble.</u>** The p-values from testing in each cell line if the validation Pearson's r is different from zero, for every transcription factor, were adjusted using the Benjamini-Hochberg correction.

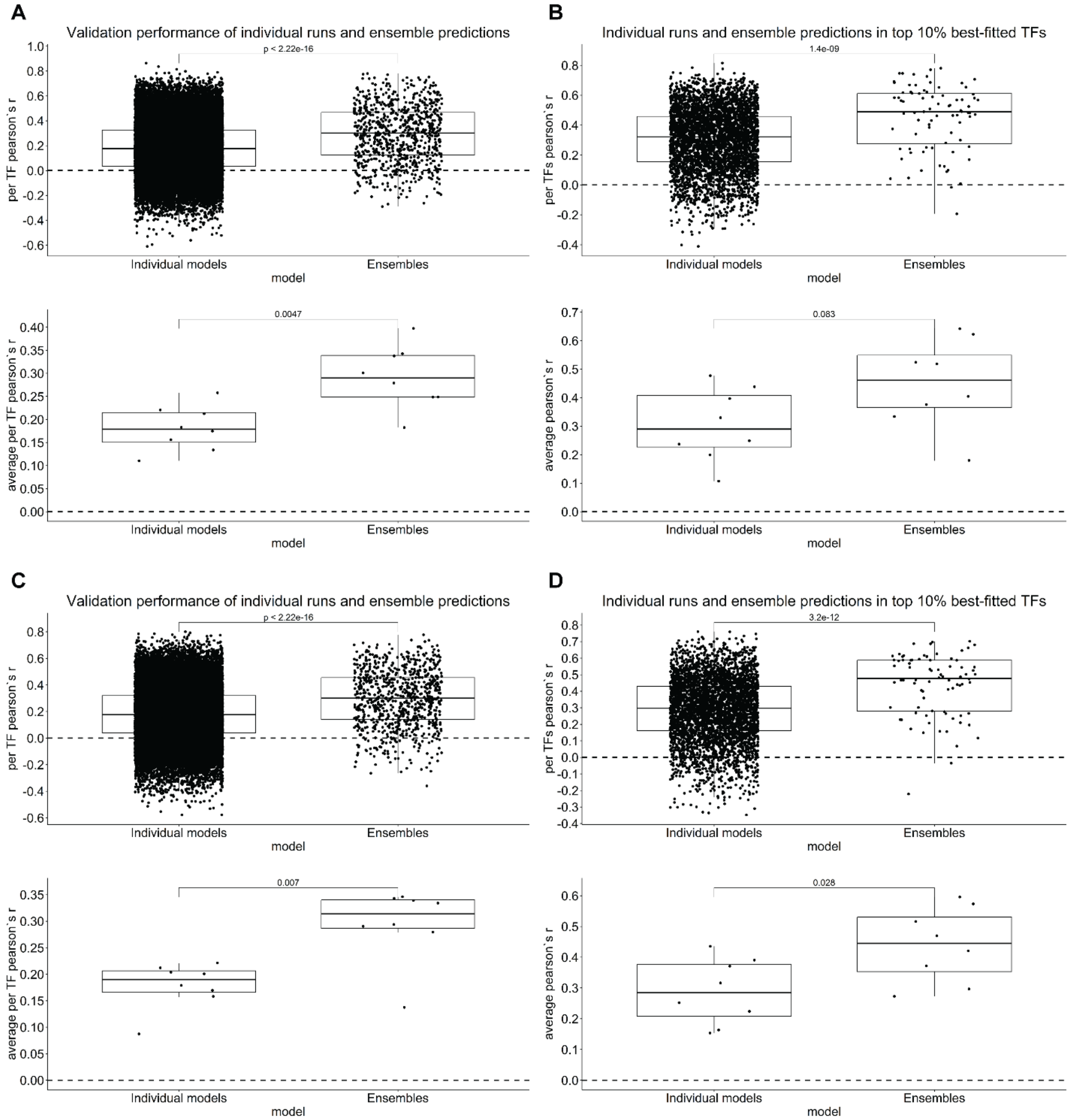


**<u>Supplementary Figure 5: Performance when using the A375 and A549 as the cell line for initial training.</u> A)** Performance of ensembles and individual models when using the A375 cell line. **B)** Performance of ensembles and individual models when using the A375 cell line, by looking only at the top 10% well-fitted TFs during training. **C)** Performance of ensembles and individual models when using the A549 cell line. **D)** Performance of ensembles and individual models when using the A549 cell line, by looking only at the top 10% well-fitted TFs during training.

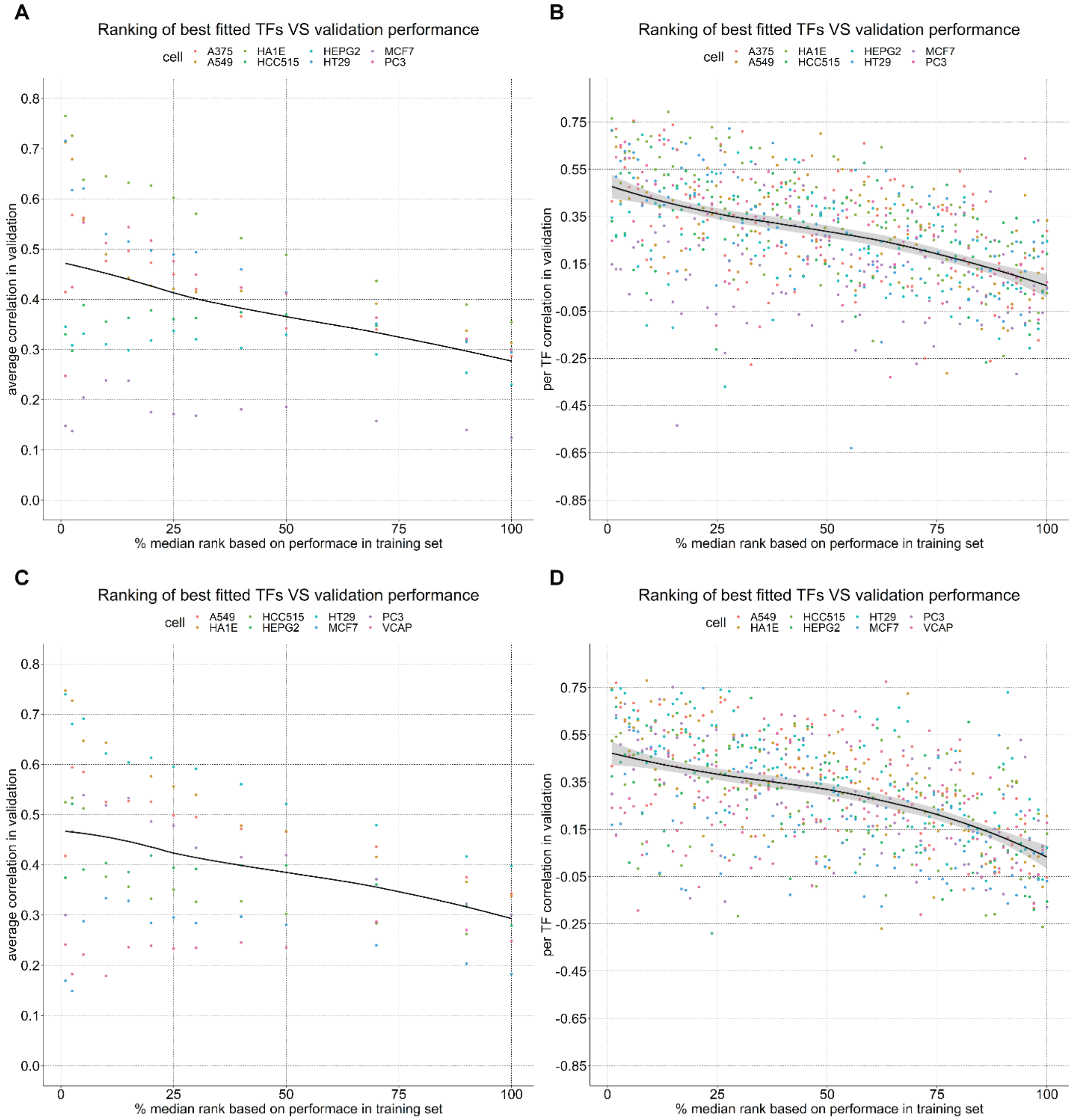


**<u>Supplementary Figure 6: Validation performance when looking only at a percentage of the best-fitted TFs during training.</u>** **A)** Average performance across all TFs for different percentage cut-offs of training performance, when using the VCAP cell line for initial training. **B)** Performance per TF for different percentage cut-offs of training performance, when using the VCAP cell line for initial training. **C)** Average performance across all TFs for different percentage cut-offs of training performance, when using the A375 cell line for initial training. **D)** Performance per TF for different percentage cut-offs of training performance, when using the A375 cell line for initial training.

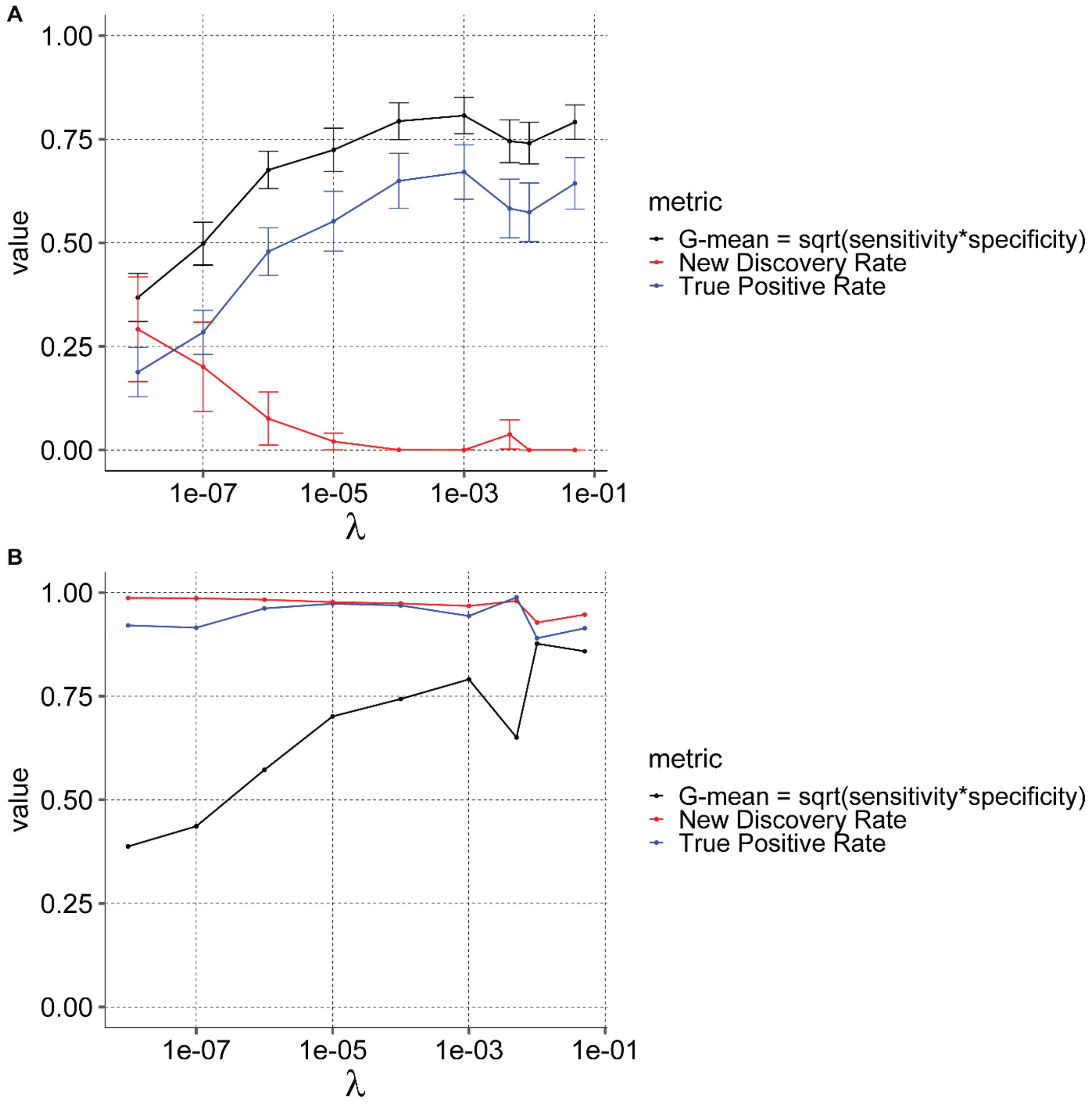


**Supplementary Figure 7: Inference of drug-target interactions is affected by regularization in VCAP cell line A)** The geometric mean of sensitivity and specificity (G-mean) when inferring new drug-target interactions at different levels of regularization. The error bars denote one Standard Error (SE) from the mean. The shaded area displays the 95% confidence interval around the smooth fitted line. **B)** The geometric mean of sensitivity and specificity (G-mean) and the NDR for the same gradient cut-off, when inferring new drug-target interactions at different levels of regularization, while using the error-based method discussed in the main manuscript.

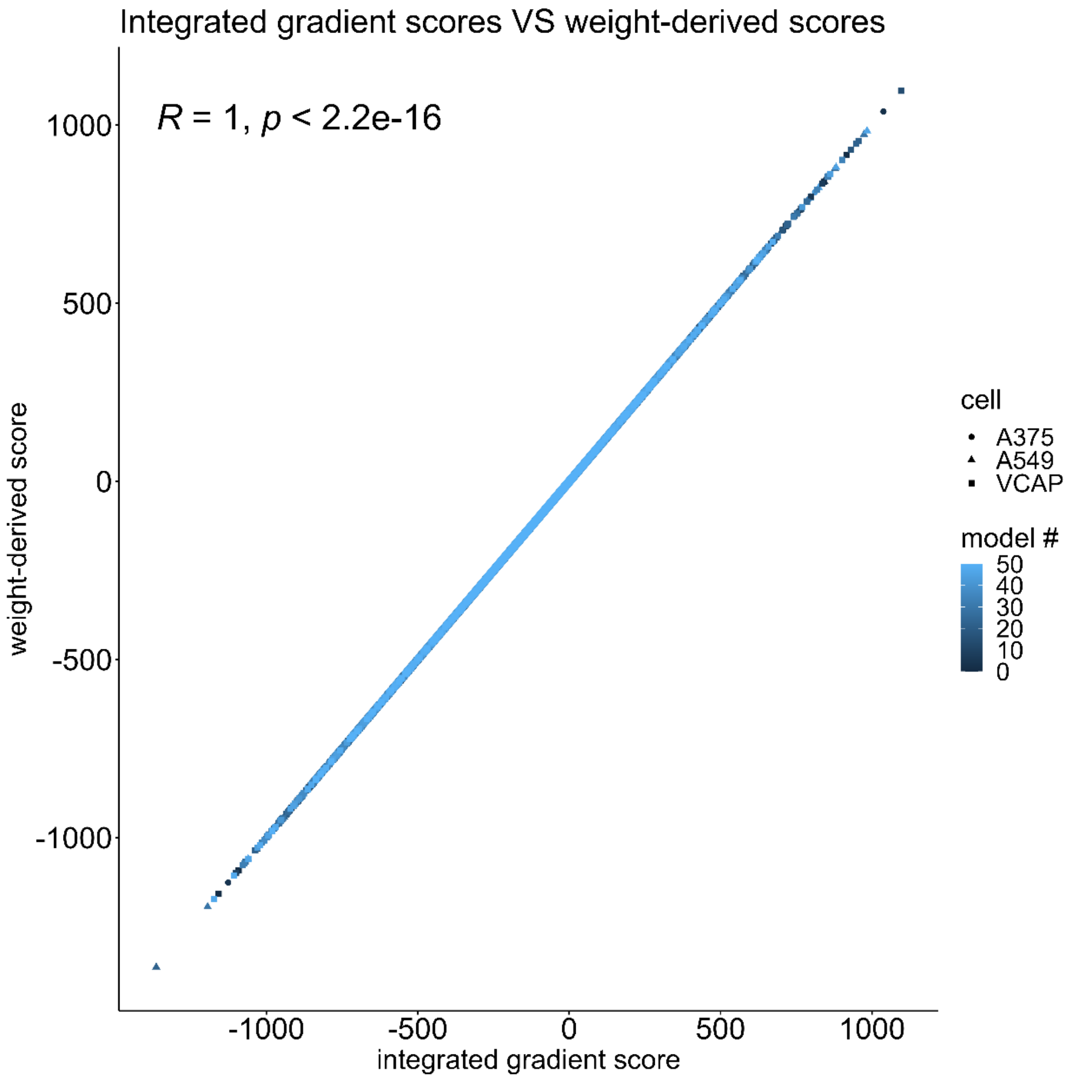


**Supplementary Figure 8:** Relationship between the integrated gradient score[1], generated from the Captum library[2], and the dug-target interaction weight score, derived from all the linear algebra operations in the linear drug layer.

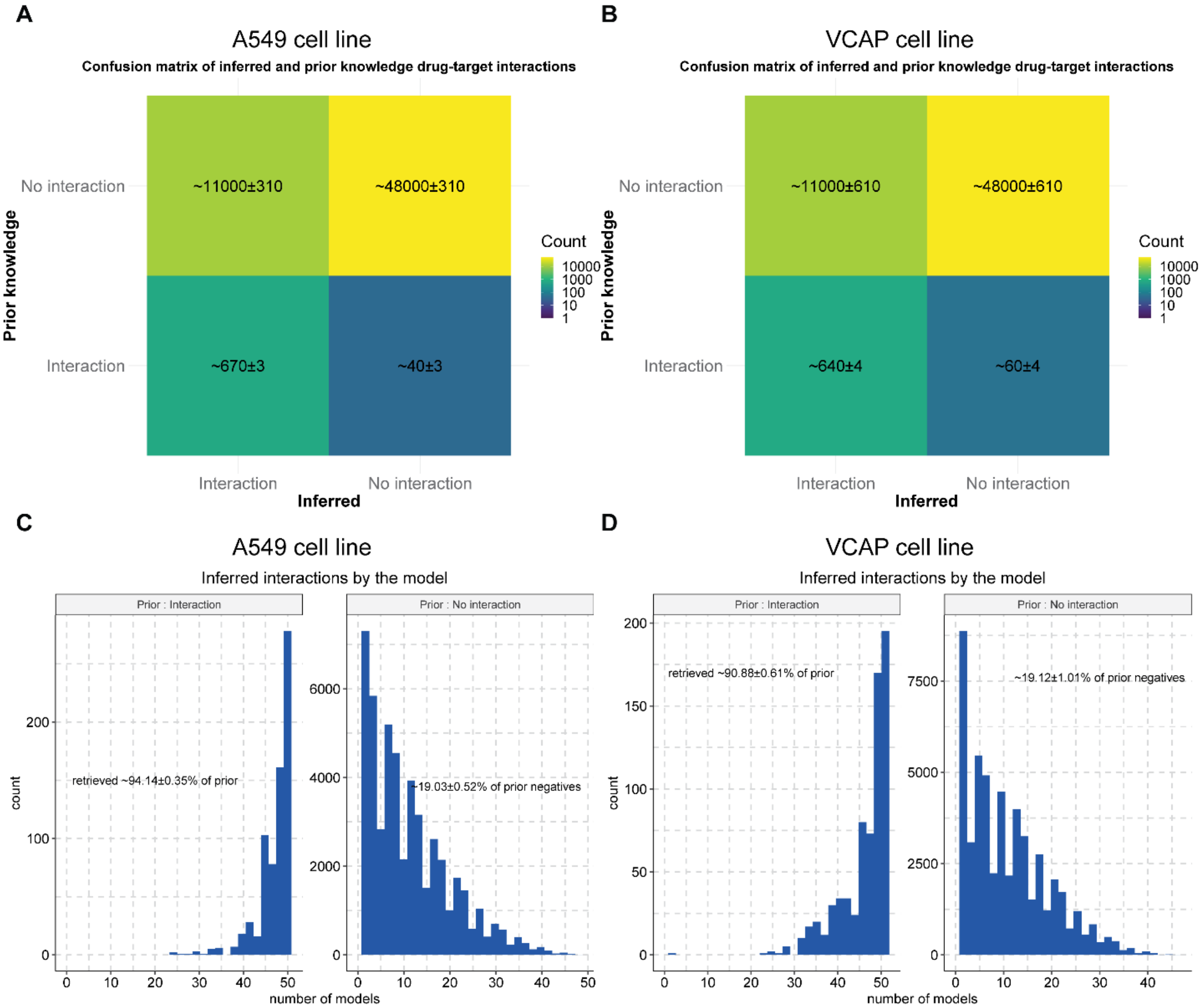


<u>**Supplementary Figure 9: Inferred drug-target interactions using the drug layer.**</u> **A-B)** Average confusion matrix using multiple trained models for the inferred drug-target interactions. **C-D)** Percentage of prior knowledge of drug-target interactions and previously unknown interactions retrieved, and their corresponding frequency of appearance in multiple models.

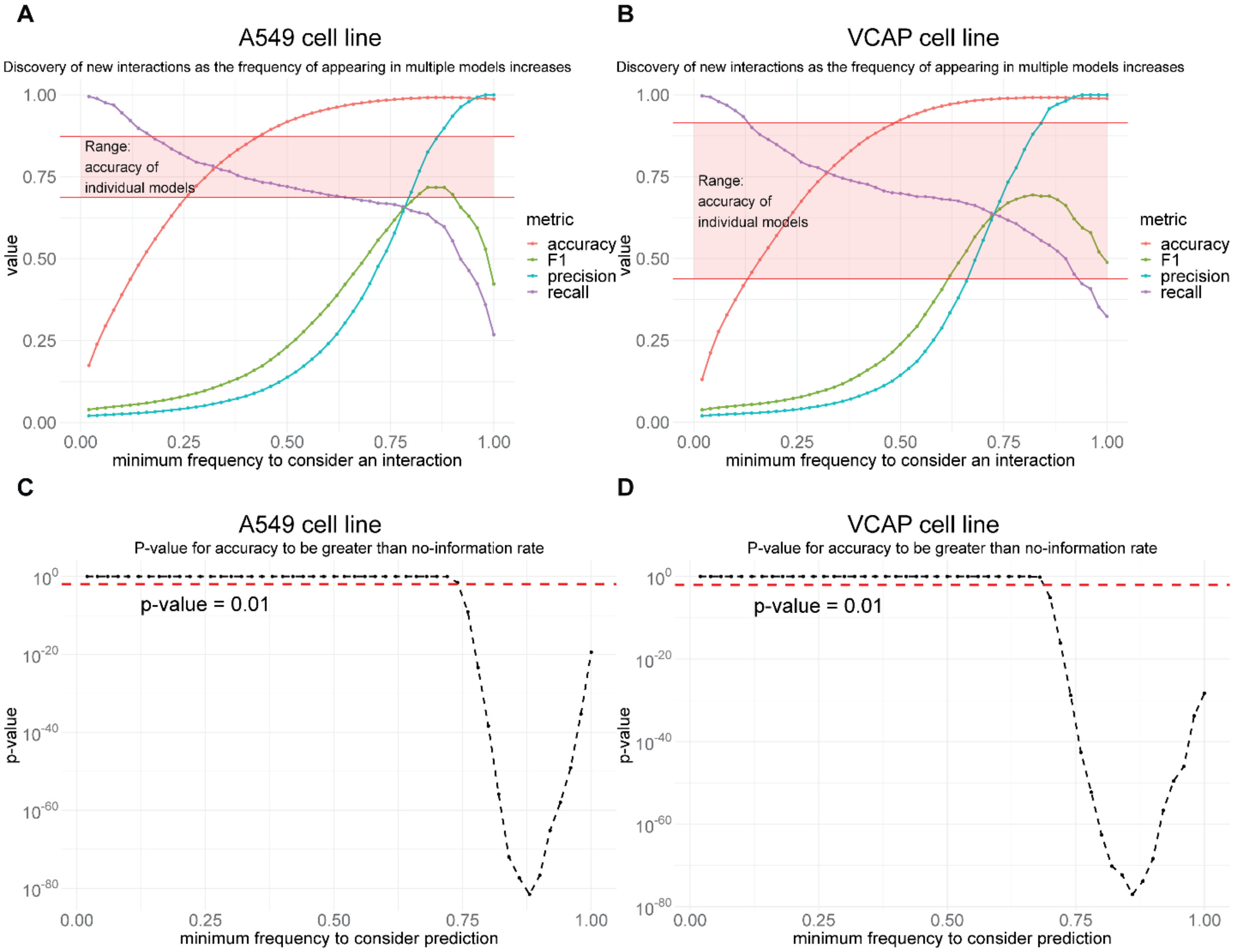


**Supplementary Figure 10: Inference analysis of drug-target interactions appearing in multiple models.** **A-B)** Classification performance of our approach by considering as ground truth the interactions contained both in the Broad's Institute Repurposing Hub[3] and in DrugBank[4]. **C-D)** P-values from comparing accuracy with the accuracy obtained by assigning everything to the predominant class (No Information Rate: NIR), for multiple frequency scores

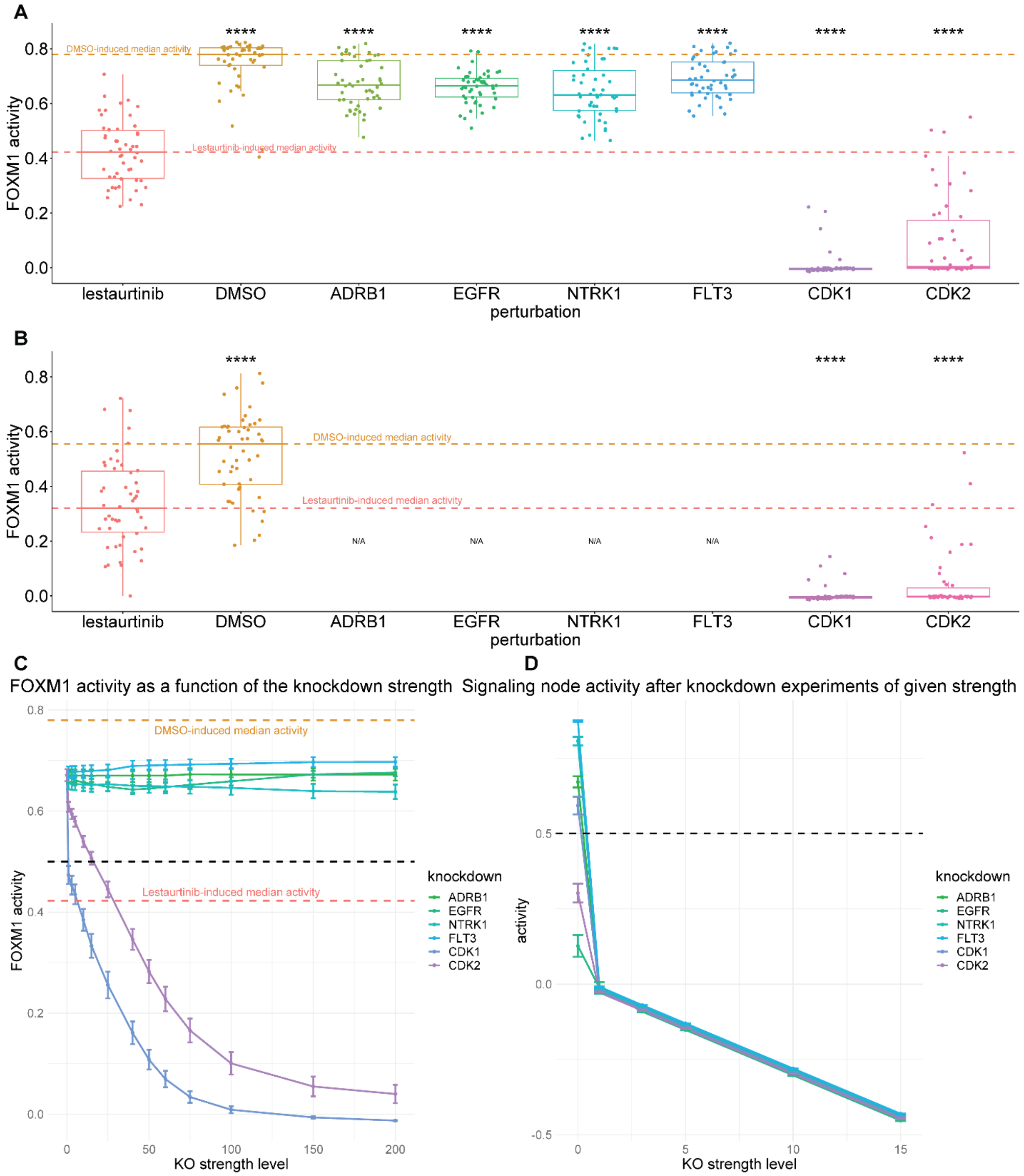


**<u>Supplementary Figure 11: FOXM1 activity after *in-silico* knockdown perturbations</u>**: **A)** *In-silico* knockdown using the trained models in the A375 cell line, with a KO strength of -100. **B)** *In-silico* knockdown using the smaller inferred network explaining the MoA of the off-target effect of Lestaurtinib in the A375 cell line, with a KO strength of -100. **C)** FOXM1 activity (using the full models) for an increasing strength of KOs. **D)** Knocked-down nodes' activity for an increasing strength of KOs (using the full models).

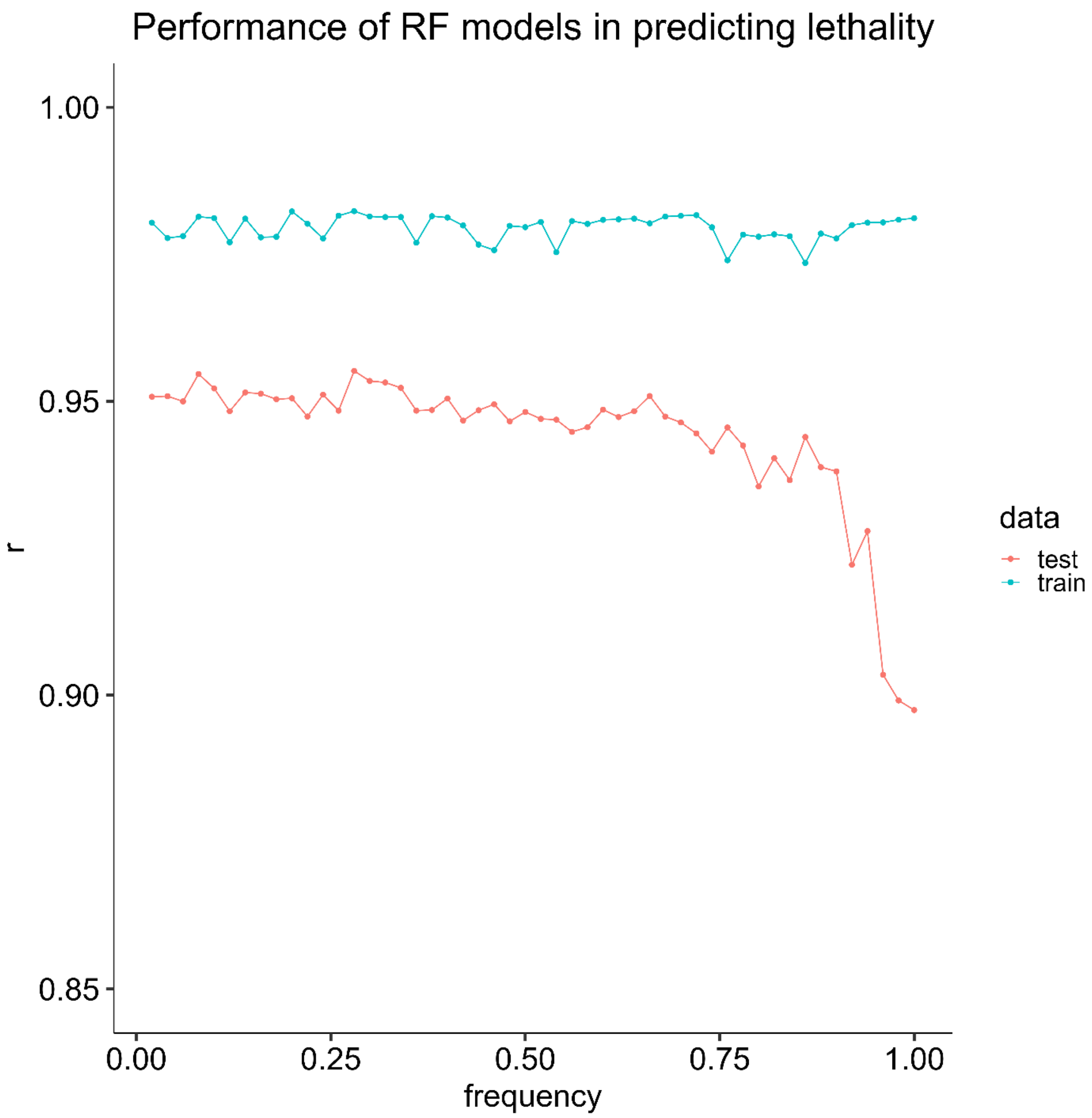


**Supplementary Figure 12: Performance of RF models in predicting lethality using inferred interactions, considered by multiple models.**

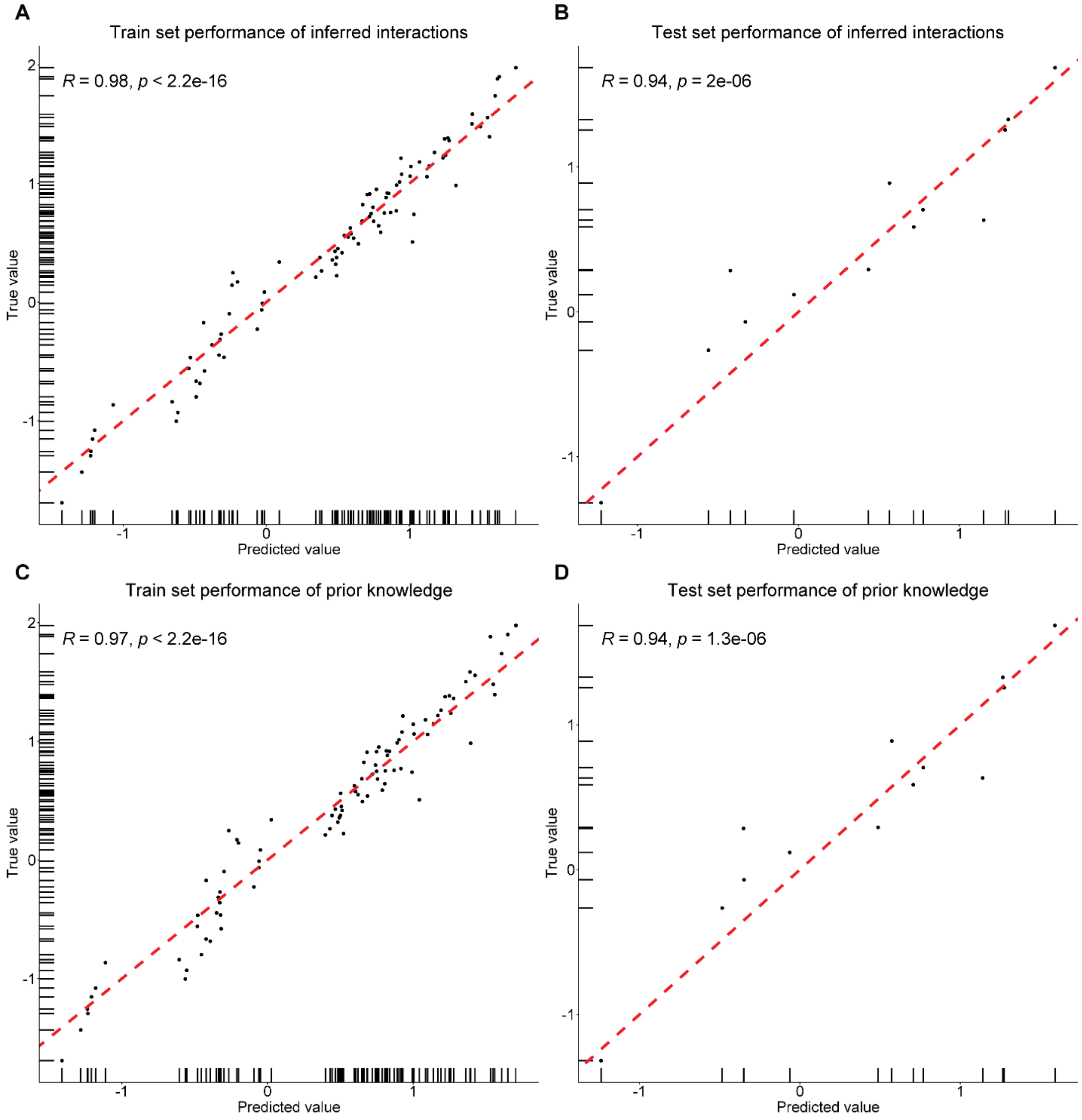


**Supplementary Figure 13: Comparison of random forest (RF) models using inferred or only known interactions to predict viability**. **A-B)** Performance of RF model trained to predict lethality from the inferred drug-target interactions, which are found by the ensemble approach if they appear in at least 44/50 models. **C-D)** Performance of RF model trained to predict lethality using only prior knowledge of drug-target interactions.

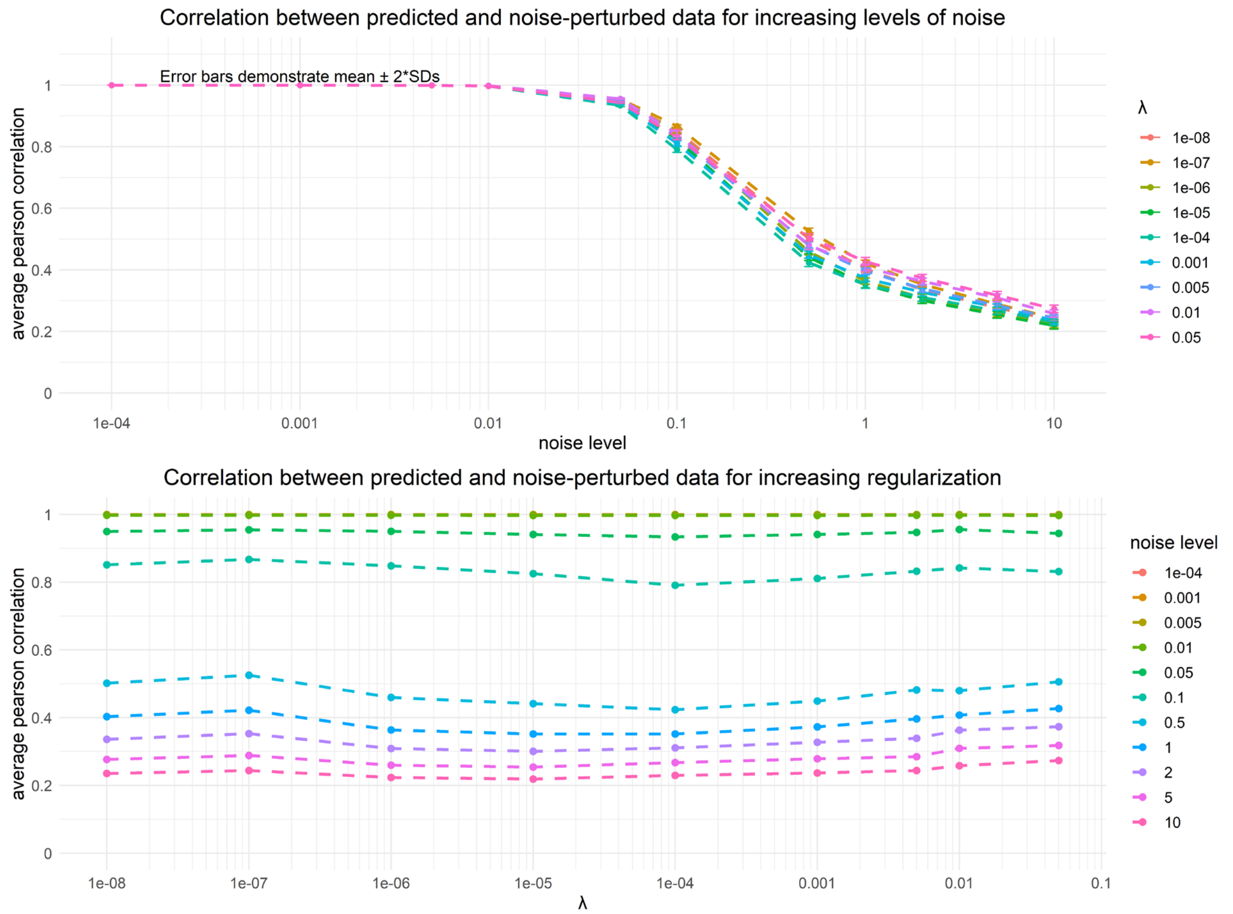


**Supplementary Figure 14: Correlation between model prediction and model prediction when adding noise in the input of LEMBAS.** The correlation is presented as a function of the drug layer regularization and the level of the noise.

### Supplementary Tables

**Table 1:** Hyper-parameters for training a whole model

| Parameter | Value |
|---|---|
| # of drugs | 233 |
| # of targets | 259 |
| Use precalculated ECFP4 similarity | TRUE |
| # of signaling nodes | 2059 |
| # of edges in the signaling net | 12127 |
| # of transcription factors | 101 |
| LEMBAS iteration steps | 120 |
| Leak (for leaky ReLU) | 0.01 |
| epochs | 5000 |
| Batch size | 25 |
| optimizer | Adam |
| Maximum Learning rate | $2*10^{-3}$ |
| Minimum Learning rate | $10^{-8}$ |
| Weight decay | 0 |
| Noise Level | 10 |
| L2 regularization | $10^{-6}$ |
| Spectral regularization | $10^{-3}$ |
| State regularization | $10^{-5}$ |
| Projection L2 regularization | $10^{-6}$ |
| Sign violation regularization | 0.1 |
| Target precision for spectral loss | $10^{-6}$ |
| Exponential factor for spectral loss | 10 |
| inputAmplitude | 3 |
| projectionAmplitude | 1.2 |

### Supplementary Notes

#### Supplementary Note 1: Regularization terms of the signaling network

We incorporated auxiliary terms in the training loss of the model from the GitHub repository (https://github.com/Lauffenburger-Lab/LEMBAS) of the LEMBAS framework[5], in order to constrain different aspects of the signaling network. First of all, for the signaling network part of the model we want to constrain the model in biologically feasible solutions, thus the learned weights need to have the same sign as the known sign of protein-protein interaction. This is done by introducing a loss heavily penalizing the violation of known signs: $signConstraint = 0.1 * \sum_{i=1}^{V} |w_i|$, where $V$ is the total number of violations and $w_i$ is a weight in the network. To prevent the fitting of parameters with extreme values, L2 regularization of the weights ($NetWeightLoss$) and biases ($biasLoss$) of the intracellular signaling network was implemented by adding the sum of squares of these vectors multiplied by $10^{-6}$. Additionally, to prevent weights from getting stuck at zero an additional term was added forming the final regularization term of the signaling weights as: $NetWeightLoss = 10^{-6} * \sum\left(w_i^2 + \frac{1}{w_i^2+0.5}\right)$. Furthermore, the trainable weights used to project from the signaling state to TF activity were also L2-regularized to avoid extreme values: $projectionLoss = 10^{-6}\sum\left(w_{pi} - 1.2\right)^2$. To ensure a dynamic range of signaling states for the signaling nodes in the intracellular network, we regularized the state variables so that each one of them has a uniform distribution across conditions, and this was implemented by regularizing some of the statistical properties to match the corresponding properties of a uniform distribution on the interval [0,0.99]. The regularization was implemented by calculating the deviation of the empirical properties of the distribution (mean, variance, maximum, and minimum value) across conditions from the ideal property calculated for the given interval, using the sum of squared errors. Additionally, as already described in the methods section, the model was penalized with a factor of 10, when the maximum value of the signaling states was negative, and finally, all contributions were added into one term ($stateLoss$) and scaled in the total loss with a coefficient of $10^{-5}$. Finally, following the implementation proposed in the LEMBAS framework[5], to ensure that the model achieves convergence by reaching a steady state we aim to constrain the absolute value of the largest eigenvalue of the transition matrix, i.e., the spectral radius (ρ), to be less than 1. This is implemented with an exponential barrier function, used to constrain the spectral radius (ρ) where: $spectralRadiusLoss = \frac{1}{e^{10*[target\ \rho]}} * (e^{10*\rho} - 1), [target\ \rho] = e^{\frac{\ln(10^{-6})}{120}}$.

**<u>Supplementary Note 2:</u> Node and edge importance for regulating the activity of a transcription factor in the signaling network**

The gradient of the nodes' biases ($db$) quantifies how much the node can affect the TF and the gradient of the weights ($dw$) how important an edge is. However, these just describe how much the activity of a TF will change if an edge, or a node, changes. To account for the importance of an edge in the current trained state we use a score combining the current weight and the gradient score: $score_w = |dw| * |weight|$. For nodes, we want to also account for how sensitive they are to changes in the signal, so we combine the node's maximum range for different signal strengths and the gradient score: $score_b = |db| * |range|$, where $range = \max(X_{in}[:, node]) - \min(X_{in}[:, node])$.

#### References


1. Sundararajan, M., Taly, A. & Yan, Q. Axiomatic Attribution for Deep Networks. in *Proceedings of the 34th International Conference on Machine Learning* 3319–3328 (PMLR, 2017).

2. Kokhlikyan, N. *et al.* Captum: A unified and generic model interpretability library for PyTorch. Preprint at https://doi.org/10.48550/arXiv.2009.07896 (2020).
3. Corsello, S. M. *et al.* The Drug Repurposing Hub: a next-generation drug library and information resource. *Nat Med* **23**, 405–408 (2017).
4. Wishart, D. S. *et al.* DrugBank: a comprehensive resource for in silico drug discovery and exploration. *Nucleic Acids Research* **34**, D668–D672 (2006).
5. Nilsson, A., Peters, J. M., Bryson, B. & Lauffenburger, D. A. *Artificial neural networks enable genome-scale simulations of intracellular signaling*. 2021.09.24.461703 https://www.biorxiv.org/content/10.1101/2021.09.24.461703v1 (2021) doi:10.1101/2021.09.24.461703.
